# An anaphase switch in astral microtubule dynamics specifically requires the APC/C^Cdc20^-dependent degradation of the mitotic cyclin Clb4

**DOI:** 10.1101/2022.04.21.470331

**Authors:** Federico Zucca, Jiaming Li, Clara Visintin, Steven P. Gygi, Rosella Visintin

**Affiliations:** Department of Experimental Oncology, European Institute of Oncology IRCCS, via Adamello 16, Milan 20139, Italy; Department of Cell Biology, Harvard Medical School, 240 Longwood Avenue, Boston, Massachusetts 02115, USA

## Abstract

Key for accurate chromosome partitioning to the offspring is the ability of mitotic spindle microtubules to respond to different molecular signals and remodel their dynamics accordingly. Spindle microtubules are conventionally divided into three classes: kinetochore, interpolar and astral microtubules (kMTs, iMTs and aMTs, respectively), among all aMT regulation remains elusive. Here, we show that aMT dynamics are tightly regulated. aMTs remain unstable up to metaphase and are stabilized at anaphase onset. This switch in aMT dynamics, crucial for proper spindle orientation, specifically requires the degradation of the mitotic cyclin Clb4 by the Anaphase Promoting Complex bound to its activator subunit Cdc20 (APC/C^Cdc20^). These data highlight a unique role for mitotic cyclin Clb4, provide a framework to understand aMT regulation in vertebrates and uncover mechanistic principles of how the APC/C^Cdc20^ choreographs the timing of late mitotic events by sequentially impacting on the three classes of spindle microtubules.

## Introduction

Chromosome segregation requires remodeling of the mitotic spindle. The mitotic spindle is composed of microtubules, Microtubule Associated Proteins (MAPs) and motor proteins [1]. Based on their function, microtubules are divided into three categories [2]: (i) kinetochore microtubules (kMTs), which connect the spindle poles to chromosomes and direct their segregation; (ii) interpolar microtubules (iMTs), which form a bundle of antiparallel microtubules and promote the distancing of the two sister chromatids *via* spindle elongation; and (iii) astral microtubules (aMTs), which connect the spindle poles to the cellular cortex and guide chromosomes along the polarity axis by dictating spindle positioning and orientation. Proper spindle positioning is not only fundamental for the correct segregation of chromosomes, but it is also crucial for many cellular processes such as stem cell maintenance, tissue homeostasis and development [3]. Despite their fundamental role, the molecular mechanisms that regulate aMT dynamics remain elusive and largely overlooked. Here we investigate aMT dynamics in *Saccharomyces cerevisiae* and whether and how they are coordinated with other cell cycle events.

Chromosome segregation is initiated by the activation of the Anaphase Promoting Complex or Cyclosome (APC/C) in complex with its activator subunit Cdc20 [4], [5]. The APC/C is an E3-ubiquitin ligase whose specificity is dictated by the interaction with its regulatory subunits Cdc20 and Cdh1 [6]. Activation of the APC/C^Cdc20^ at the metaphase to anaphase transition initiates a three-step signaling cascade that culminates in cohesin cleavage - the point of “no-return” for mitotic exit. Cohesin is a protein complex that holds chromosomes together from the moment of their replication up to their separation [7]. Following cohesin cleavage, the coordination between sister chromatid separation and segregation is directed by changes of mitotic spindle microtubule dynamics. Cohesin cleavage indirectly affects the dynamics of nuclear microtubules – namely kMTs and iMTs – by removing the opposing forces to spindle pulling. kMTs retract and pull sister chromatids toward the spindle poles while iMTs drive spindle elongation, thereby segregating sister chromatids apart from each other. Consistent with their functions (kMTs search and capture chromosomes and iMTs promoting the formation of a short bipolar spindle without forcing elongation), nuclear microtubules are unstable in metaphase [8]. *Vice versa,* to preserve proper kMT-chromosome interactions and to promote spindle elongation kMTs and iMTs are stabilized in anaphase [8], [9]. From a molecular point of view, this switch in microtubule dynamics from an unstable to a stable state has long been associated with Cyclin Dependent Kinase (CDK) activity. CDK activity is high in metaphase and decreases in anaphase due to APC/C^Cdc20^-mediated cyclin degradation and activation of the main yeast CDK-counteracting phosphatase Cdc14. A long-standing model, supported by the observation that several motor and microtubule associated proteins (MAPs) are regulated by phosphorylation and de-phosphorylation events mediated by the antagonistic couple CDK1/Cdc14 [10]–[12], foresaw that spindle microtubules were stabilized because of an overall change in the phosphorylation landscape in favor of de-phosphorylation. This view has been recently challenged by an elegant study in budding yeast that reported that the number of phosphorylated or de-phosphorylated residues in late mitosis is similar [13], thus suggesting that mitotic kinases can compensate for the drop in CDK-mediated phosphorylation. Supporting this model is the observation that the phosphatase Cdc14 and the polo-like kinase Cdc5 are redundant in triggering spindle elongation [14].

The contribution of spatially-defined mechanisms renders the dynamics of microtubule regulation more complex. An example is the phospho-regulation of single kMTs following their binding to kinetochores, which relies on the inhibition of the Aurora B kinase to promote kMT stabilization when the attachment that is limited to a single kinetochore creates tension [15], [16]. Altogether, these observations unveiled a sophisticated regulation of nuclear spindle microtubules taking place at the metaphase to anaphase transition, which relies both on phosphorylation and de-phosphorylation events – depending on the residue –, and takes into account the spatially defined modulation of individual microtubules. How aMTs fit into this picture remains unknown. Up to date, aMTs regulation has been tackled by investigating how binding to the cellular cortex alters their dynamics. It emerged that, both in budding yeast and higher eukaryotes, CDKs negatively affect this interaction. In yeast, the cyclin Clb4 promotes the detachment of aMTs from the cellular cortex in early mitosis[17] by phosphorylating a yet-to-define substrate. In human, CDK1 reduces aMTs binding to the membrane by phosphorylating nuclear mitotic apparatus (NuMa)[18]. NuMa localizes at the cortex and it is a component of the evolutionary conserved cortical machinery essential for spindle orientation [19]. The human Polo-like kinase 1 (Plk1) phosphorylates NuMa[20] and contributes to negatively regulating its cortex localization [21]. Instead, little is known as to whether astral microtubules dynamics are regulated in a cell cycle dependent manner. The only evidence of a direct regulation of aMT dynamics comes from a recent study [22], highlighting a CDK1-dependent phosphorylation of the plus-end tracking protein GTSE1 required to destabilize aMTs in prometaphase. Here we show that, similarly to iMT and kMT microtubules, aMTs dynamics switch from an unstable to a more stable status at the metaphase to anaphase transition and that at the heart of this switch - required to establish and maintaining proper aMT-cellular cortex interactions and ultimately proper spindle positioning - is the APC/C^Cdc20^-dependent degradation of the mitotic cyclin Clb4. Besides isolating the unique role of Clb4, among all cyclins, in this process, our data allow us to put forward a model that envisions the APC/C^Cdc20^ as the master choreographer of late mitotic events by sequentially instructing the three classes of spindle microtubules.

## Results

### Astral microtubules are stabilized in anaphase

To gain insights into astral microtubule (aMT) regulation in mitosis, we probed aMT morphology in *Saccharomyces cerevisiae* during an unperturbed cell cycle. To this aim, wild-type cells were synchronously released from a G1 block and cell cycle progression was assessed by monitoring nuclear and spindle morphologies (DAPI and Tub1 staining, respectively). aMT length and number, two *bona-fide* indicators of aMT stability [23], were measured at each time-point. A correlation emerged between the population approaching metaphase and a decrease in both aMT length and number (**figure 1A**). Contrariwise, these parameters increased when the population entered anaphase (**figure 1A**). This relationship became particularly evident when we correlated the morphology of aMTs with metaphase and anaphase cells (going from a mean of 2µm and 1.6 aMT/cell in metaphase cells to 2.7µm and 3 aMT/cell in anaphase cells; **figure 1B**). Of note, in budding yeast, metaphase cells have short bipolar spindles (1.5–2μm) and undivided nuclei, while anaphase cells carry elongated spindles (up to 10μm during telophase) and have separated DNA masses (**figure 1C**).

**Figure 1.**
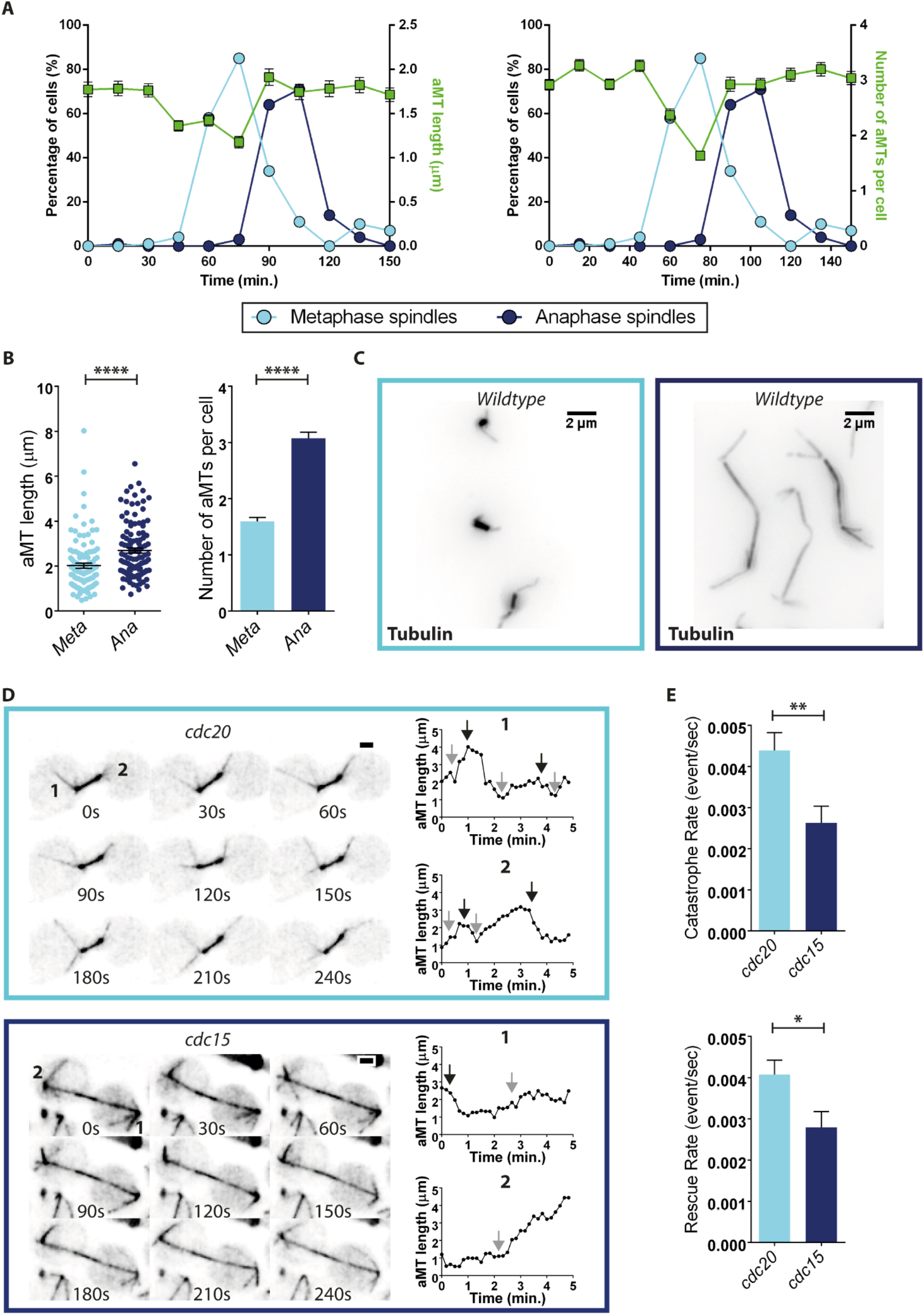
aMTs are stabilized in anaphase. (A-C) Wild-type (Ry1) cells were synchronized in G1 with α-factor (5µg/ml) and released into fresh YEPD media lacking the pheromone. (A) At the indicated time points, the percentage of cells containing metaphase (light blue circles) and anaphase (dark blue circles) spindles was determined and aMT length and number were measured (n=100 cells). (B) aMT length and number were measured in metaphase (60 minutes time point) or anaphase (100 minutes time point) cells (n=100, ****=p<0.0001; * asterisks denote significant differences according to unpaired t-test). (C) Representative images of wild-type metaphase (light blue) and anaphase (dark blue) spindles are shown, scale bar = 2µm. NOTE: Here and throughout the manuscript, aMT length is shown in dot plots while aMT number is shown in histograms. Each data point in the dot plot represents one single aMT. The median is displayed as a solid line. Error bars represent the Standard Error of the Mean (SEM). (D-E) *cdc20-AID* (Ry7732) and *cdc15-as1* (Ry9741) cells harboring a *TUB1-GFP* fusion were arrested in G1 and synchronously released in flask at 37°C, into fresh media supplemented with Auxin (500µM) and the 1NM-PP1 analogue 9 (5µM) to inactivate the *cdc20-AID* and *cdc15-as1* alleles, respectively. When the majority of the cells reached their terminal arrest (∼3 hours after the release), cells were moved to a CellASIC ONIX plate and z-stack time-lapse images of spindle microtubules were acquired every 10 seconds for 5 minutes to determine the length of individual aMTs. (D) Time-lapse images of a representative cell are shown for both *cdc20-AID* and *cdc15-as1* strains (scale bar = 1µm). Graphs of two representative aMTs (1 and 2) *per* mutant are shown. Black and gray arrows indicate the occurrence of catastrophe and rescue events, respectively. (E) Histograms represent the calculated catastrophe and rescue rates (n=33 aMTs in *cdc20-AID* cells and n=21 aMTs in *cdc15-as1* cells were measured; *=p<0.1; **=p<0.01, * asterisks denote significant differences according to unpaired t-test).

Since the distinction between cells in late G2 and in early metaphase, or in late metaphase and in early anaphase, is somewhat arbitrary, we moved to probe aMT dynamics in homogenous populations and assessed aMT morphology in *cdc20* and *cdc15* mutant cells, which arrest in metaphase and anaphase, due to impairment in the activation of APC/C^Cdc20^ and mitotic exit network [24], respectively. To arrest cells in metaphase, unless otherwise specified, we used a conditional allele of *CDC20* where the wild-type *CDC20* gene is fused to an Auxin-inducible degron sequence (*CDC20-AID* henceforth *cdc20).* This allele encodes for a protein that is selectively degraded in the presence of auxin [25]. To arrest cells in anaphase, we used an ATP analog-sensitive allele of the kinase *CDC15* (*cdc15-as1*) that is inactivated upon addition of the Cdc15-as1 inhibitor 1NM-PP1[26] to the cell culture media. *cdc20* and *cdc15* mutants were released from a G1 block into the restrictive conditions for the alleles, and aMTs were analyzed when cells reached their terminal phenotype (approximately 3 hours after the release, as assessed by spindle morphology). While the arrest uniformed the average aMT length between metaphase and anaphase cells (2.3µm in *cdc20* cells and 2.1µm in *cdc15* cells), the number of aMTs remained significantly higher in *cdc15* mutants (from a mean of 3.1 to 4.9 aMT/cell in *cdc20* and *cdc15* cells, respectively; **supplementary figure 1**). To exclude possible mutant-specific effects, the same assays were performed in additional mutants that arrest either in metaphase (*cdc20-1* and *cdc23-1* cells, Cdc23 is a core subunit of APC/C [27]) or in anaphase (*cdc5-as1* [28] and *cdc14-1* [29] cells). The data obtained confirmed what we found in *CDC20-AID* and *cdc15-as1* cells (**supplementary figure 1**), thus indicating that changes in aMT morphology are dictated by the cell cycle phase rather than being a specific trait of the mutant strains.

The observation that upon protracted arrests the difference in aMT length flattened between the two cell cycle phases prompted us to explore aMT dynamics. For this purpose, we moved from fixed-cell samples to live cell-imaging [30], an approach that allows us to extract parameters that ultimately define dynamic properties, such as catastrophe and rescues rates [31], [32]. The length of individual aMTs was measured over time in single cells and probed in *cdc20* and *cdc15* cells expressing a GFP-tagged Tub1 fusion protein at their terminal phenotype (**figure 1D**). In line with previous findings (**supplementary figure 1**), aMTs of *cdc15* cells resulted less dynamic - as assessed by the decrease of both catastrophe (from 0.0044 to 0.0026 event/sec) and rescue rates (from 0.004 to 0.0028 event/sec) - than aMTs of *cdc20* cells (**figure 1E**). Taken together, the observations that: (i) aMTs increase in number and length when wild-type cells move from metaphase to anaphase; (ii) aMT number is higher in mutants arrested in anaphase when compared to their metaphase counterpart; and (iii) anaphase aMTs appear less dynamic, support the conclusion that aMTs are stabilized at anaphase onset. Why aMT length is similar in the two conditions remains puzzling. However, it is possible that while the length of metaphase aMTs at steady state is determined solely by their catastrophe and rescue rates, the length of anaphase aMTs is also negatively influenced by spindle elongation, i.e., spatial constraints due to proximity to the cellular cortex.

### Stabilization of aMTs in anaphase relies on a specific signature

Having established that, similarly to kinetochore and interpolar microtubules, aMTs as well are unstable in metaphase and stabilized at anaphase onset, we wished to characterize the molecular mechanism at the heart of this switch. Although it remains unclear how the anaphase stabilization of kMTs is achieved, it is known that the activity of either the phosphatase Cdc14 or the Polo-like kinase Cdc5 [14] is essential for anaphase stabilization of iMTs. To investigate whether astral and interpolar microtubules share the same regulatory mechanism, we examined the aMT phenotype of *cdc14 cdc5* double mutant cells [14]. These cells arrest in mini-anaphase (after cohesin cleavage, the conventional indicator of anaphase onset), but carry short bipolar spindles and undivided nuclei, which is typical of metaphase cells [14], due to defects in spindle elongation. Although it was reported that *cdc14 cdc5* double mutants do not elicit any significant checkpoint response [14], in our characterization of the aMT phenotype in these cells, we also abrogated the checkpoint responses by deleting *MAD2*, *RAD9* and *BUB2*, which code for essential components of the Spindle Assembly-, the DNA Damage- and the Spindle Positioning-Checkpoint, respectively. Building on the knowledge that aMT dynamics are dictated by the cell cycle phase and that while the SAC and the DDC arrest cells in metaphase, the SpoC acts and arrests cells later in anaphase, we combined *MAD2* and *RAD9* deletions but kept them separate from the deletion of *BUB2*.

aMT dynamics of *cdc14 cdc5* cells were probed at their terminal phenotype both by indirect immunofluorescence and live-cell imaging, and compared with the ones of *cdc20* and *cdc15* arrested cells, which have similar spindle morphology and mitotic phase, respectively, of the double mutant cells. Consistent with the anaphase arrest of *cdc14 cdc5* cells, aMTs of these cells were significantly more stable than aMTs of metaphase-arrested *cdc20* cells: they were both longer and more numerous (from a mean of 2.6µm and 2.6 aMT/cell in *cdc20* cells to a mean of 4µm and 4.6 aMT/cell in *cdc14 cdc5* cells; **figure 2A** and **2B**). This result suggests that the molecular mechanism at the core of the anaphase aMT stabilization differs from the one controlling iMTs, as it is independent of Cdc14 and/or Cdc5 activities and implies that spatial constraints may play a role.

**Figure 2.**
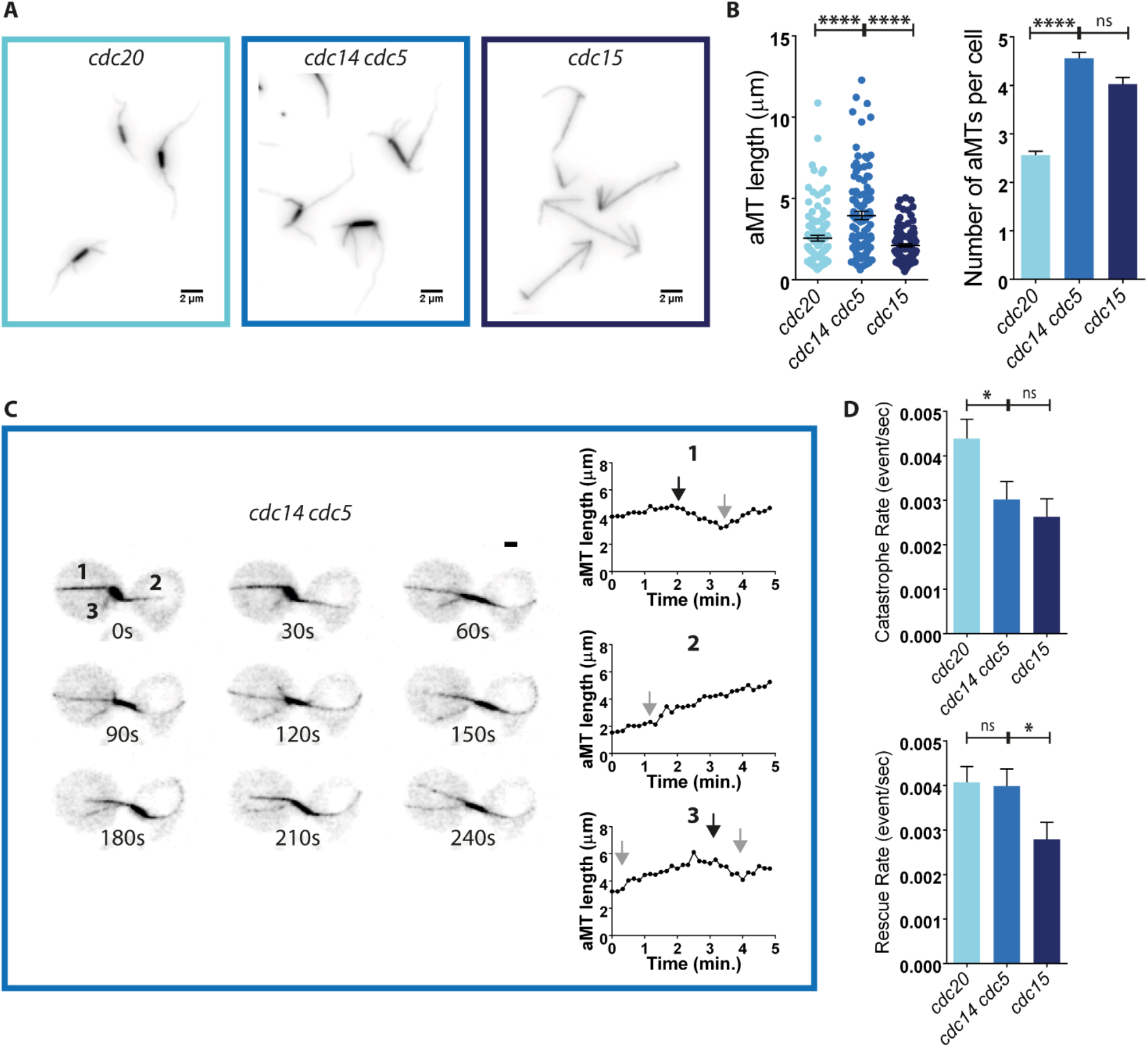
aMT stabilization relies on a specific anaphase signature. (A-B) *cdc20-AID* (Ry4853)*, cdc14-1 cdc5-as1* (Ry1602) and *cdc15-as1* (Ry1112) cells were arrested in G1 with α-factor (5µg/ml) and synchronously released at 37°C, to inactivate the *cdc14-1* allele, into fresh YEPD media, supplemented with Auxin (500µM), 1NM-PP1 analogue 9 (5µM) and CMK (5µM) to inactivate the *cdc20-AID*, *cdc15-as1* and *cdc5-as1* alleles, respectively. Cells were analyzed at their terminal arrest (∼3 hours after the release). (A) Representative images of *cdc20-AID, cdc14 cdc5 and cdc15* arrested cells are shown. (B) The graphs show aMT length and number of the indicated genotypes (n=100; ****=p<0.0001; * asterisks denote significant differences according to ordinary One-Way ANOVA and Tukey’s multiple comparisons test). (C-D) *cdc20-AID* (Ry7732), *cdc15-as1* (Ry9741) and *cdc14-1 cdc5-as1* (Ry3256) cells expressing *Tub1-GFP* fusion protein were treated as in (a-b). Approximately 3 hours after the release, cells were moved to a CellASIC ONIX plate and z-stack time-lapse images of spindle microtubules were acquired every 10 seconds for 5 minutes to determine the length of individual aMTs. (C) Time-lapse images of a representative cell are shown for the *cdc14-1 cdc5-as1* strain (scale bar = 1µm). Graphs of three aMTs (1, 2 and 3) are shown. The occurrence of catastrophe and rescue events is indicated in the graphs with black and gray arrows, respectively. (D) The histograms show the catastrophe and rescue rates of the indicated mutant strains (33 aMTs in *cdc20-AID* cells, 33 aMTs in *cdc14-1 cdc5-as1* cells and 21 aMTs in *cdc15-as1* cells were measured; *=p<0.1; * asterisks denote significant differences according to ordinary One-Way ANOVA and Tukey’s multiple comparisons test).

To our surprise, the aMTs of *cdc14 cdc5* mini-anaphase arrested cells were also more stable than the aMTs of late anaphase arrested *cdc15* mutants. In this case, aMTs of both *cdc14 cdc5* and *cdc15* cells were similar in number but aMTs of *cdc14 cdc5* cells were much longer than the ones of *cdc15* cells (from a mean of 2.1µm to 4µm in *cdc15* and *cdc14 cdc5* cells, respectively; **figure 2A** and **2B**). This phenotype is not caused by an inappropriate activation of checkpoint pathways since abrogation of checkpoint activities in the double mutant background did not alter aMT morphology (both *cdc14 cdc5 mad2 rad9* and *cdc14 cdc5 bub2* cells retain aMTs that are similar in length and number to aMTs of the double mutant cells - **supplementary figure 2A** and **2B**).

To understand why *cdc14 cdc5* and *cdc15* cells carry stable aMTs but of different length, we moved to live-cell imaging. It turned out that while the aMT catastrophe and rescue rates of metaphase (*cdc20*) and anaphase (*cdc15*) cells always follow the same trend, either by increasing or decreasing in number, possibly reaching an equilibrium, the two parameters are uncoupled in the aMTs of the double mutant cells (**figure 2C and 2D**). Here, the aMTs are characterized by a low-catastrophe rate (0.003 events/sec in *cdc14 cdc5* cells, compared to 0.004 and 0.0026 in *cdc20* and *cdc15* cells, respectively), which is associated with a high-rescue rate (0.004 events/sec in *cdc14 cdc5* cells, compared to 0.004 and 0.0028 in *cdc20* and *cdc15* cells, respectively; **figure 2C and 2D**), a combination that favors polymerization and could provide an explanation to the different aMT lengths observed.

Why catastrophe and rescue rates are uncoupled in *cdc14 cdc5* cells remains unclear. Different scenarios can be envisioned: (i) spindle elongation may negatively impact on astral microtubule dynamics in a direct or indirect manner (e.g., cross talk between interpolar and astral microtubules; spatial constraints imposed by bringing the poles in close proximity to the cellular cortex); (ii) Cdc14 and/or Cdc5 may directly counteract aMT stabilization; or (iii) a combination of the two. To assess the contribution of spindle elongation *per se* in aMT dynamics, we prevented the establishment of aMTs-cortex connections, in a *cdc15* background by modulating the amounts of the microtubule binding spindle positioning factor Kar9 [33] and of the minus-end directed motor Dyn1[34], two proteins implicated in the process. To prevent the synthetic lethality of combining *KAR9* and *DYN1* deletions, we performed a simultaneous inactivation of Kar9 and Dyn1 by combining the conditional *DYN1-AID* mutant allele with the *KAR9* deletion [35]. Our findings show that the aMTs of *cdc15 kar9 dyn1* cells are significantly longer than aMTs of *cdc15* cells (1.68 fold change) and resemble aMTs of *cdc5 cdc14* double mutant cells, indicating that spindle elongation *per se* is not a limiting factor in aMT stabilization (**supplementary figure 3A** and **3B**) but it rather contributes to aMT dynamics by bringing these microtubules in the vicinity of the cortex.

To assess the contribution of Cdc14 and Cdc5 in aMT regulation, we probed aMT morphology in metaphase-arrested cells lacking either Cdc14 or Cdc5 and found that aMTs of *cdc20, cdc20 cdc5* and *cdc20 cdc14* cells were similar in number and had similar lengths (**supplementary figure 3E**). To establish whether, as for iMTs, the two enzymatic activities are redundant, we combined *cdc14* and *cdc5* mutant alleles with an allele of *CDC20* whose expression is under the control of the methionine-repressible promoter (*pMET-CDC20*). We used this allelic version of *CDC20* because the concomitant inactivation of Cdc14 and Cdc5 is synthetically lethal with the tested *CDC20* mutant alleles (i.e., *cdc20-1*, *cdc20-3 and cdc20-AID*) and temperature sensitive alleles of genes encoding for core subunits of the APC/C (i.e., *cdc23)*[36], which arrest in metaphase as well. *pMET-CDC20* cells were synchronized in G1, where Cdc20 is depleted via APC/C^Cdh1^-dependent degradation, followed by a release in fresh cell culture media containing methionine to prevent *de novo* synthesis at the restrictive conditions for the alleles used in the analysis. The aMTs of *pMET-CDC20* cells carrying a mutant allele of *cdc14*, *cdc5* or both, compared to the aMTs of *pMET-CDC20* and *cdc14 cdc5* cells, exhibited an intermediate phenotype with respect to their length: the aMTs of *pMET-CDC20 cdc14*, *pMET-CDC20 cdc5* and *pMET-CDC20 cdc14 cdc5* cells were slightly longer than their metaphase counterpart (*pMET-CDC20* alone), but shorter than aMTs of the double mutant cells, and remained similar in number to aMTs of metaphase cells (**supplementary figure 3D**). Given that this intermediate phenotype was not observed in *cdc20 cdc5* and *cdc20 cdc14* cells (**supplementary figure 3E**) we believe that it is due to the less stringent metaphase arrest achieved with the *pMET-CDC20* allele (**supplementary figure 3C**). Taken together these data indicate that an anaphase-specific signature lies at the heart of the stabilization of this class of microtubules.

### APC/C^Cdc20^ activation is sufficient to stabilize aMTs at anaphase onset

Having assessed that aMT stabilization require an anaphase specific trait, we aimed to identify the molecular event that triggers this process. Although anaphase onset is conventionally defined by cohesin cleavage, this event can be dissected in three critical steps: (i) the activation of the APC/C^Cdc20^; (ii) the activation of the separase/Esp1, mediated by the APC/C^Cdc20^-dependent degradation of securin/Pds1; and (iii) the Esp1-mediated cleavage of the Scc1 subunit of the cohesin complex. Of note, each step promotes a cascade of events besides the ones involved in cohesin cleavage, for instance, Esp1 is required for the activation of the CDK-counteracting phosphatase Cdc14 [37]. Given that changes in kMT and iMT dynamics have been associated with specific cell cycle events, namely cohesin cleavage and Cdc14 activation, respectively [8], [15], [38], [39], to identify the cell cycle signal that elicits aMT stabilization, we took advantage of mutant strains that are impaired in the completion of the sequential steps of the metaphase to anaphase transition. In detail, we compared aMTs of *cdc20*, *esp1*, *scc1nc,* and *cdc14 cdc5* cells. In *cdc20* mutants, the APC/C^Cdc20^ is inactive, all its substrates are present, separase is inactive and cohesin is bound to chromatin; in *esp1* mutant cells, the APC/C^Cdc20^ is active and all its substrates are removed, including Pds1, but separase remains inactive and cohesin is still bound to chromatin; in cells expressing an uncleavable allele of Scc1 (*scc1R180DR268D)*[7], [40], henceforth *scc1nc*, not only all the APC/C^Cdc20^ substrates are removed, but also all separase substrates are properly cleaved, with the only exception of cohesin, which remains intact and bound to chromatin; and, finally, in *cdc14 cdc5* cells, all the steps of the cascade can be completed.

To assess morphology and dynamics of aMTs in the described mutants, cells were released from the G1 arrest into the restrictive conditions for the different alleles. During the analysis, we noticed that, differently from *cdc20* and *cdc14 cdc5* cells, which maintain a stable short-bipolar spindle for the entire duration of the experiment, both *esp1* and *scc1nc* cells disassembled their mitotic spindle soon after reaching a short-bipolar spindle configuration (metaphase-like morphology; **supplementary figure 4A**). This observation is consistent with data in the literature reporting that these cells can pull the nucleus, hence the spindle, into the bud and, as a consequence, exit from mitosis prematurely, albeit having failed to separate sister chromatids [41]. To perturb the system as little as possible, we initially analyzed aMTs of *esp1* and *scc1nc* cells at the time point prior to spindle disassembly, keeping in mind that we might be analyzing a mixed population of cells including some that have already reached the terminal phenotype of the mutant under investigation while others are still in metaphase. aMTs of both *esp1* and *scc1nc* cells were slightly more numerous and longer than aMTs of *cdc20* cells (from 2.3µm and 2.4 aMT/cell in *cdc20* cells to 2.9µm and 2.8 aMT/cell, and 3µm and 3.2 aMT/cell in *esp1* and *scc1nc* cells, respectively; **supplementary figure 4B** and **4C**), and they resembled the aMTs of *cdc14 cdc5* cells, hence suggesting that APC/C^Cdc20^ activation is the molecular event triggering aMT stabilization. Consistent with the risk of having a contamination of metaphase cells in this study and supporting our previous finding that homogeneous populations in protracted arrests are a better setting to accurately evaluate a phase-specific phenotype (**supplementary figure 3C-E**), the difference in aMT length and number between *cdc14 cdc5* and *cdc20* cells was less obvious when these parameters were measured at the same time point as the one chosen for *esp1* and *scc1nc* cells, namely the time point before spindle breakdown (compare **supplementary figure 4C** with **figure 2B**).

To strengthen our observation that APC/C^Cdc20^ activity is the critical event for aMT stabilization, we engineered our yeast strains to prevent spindle collapse in *esp1* mutant cells. Since spindles pulling into the bud mostly rely on the activity of the motor protein Dyn1 [32], and mitotic exit, hence spindle disassembly, is triggered by the activation of the Mitotic Exit Network (MEN)[41] in the bud, which ultimately leads to the activation of Cdc14, we decided to: (i) prevent spindle movement into the bud by disrupting Dyn1-mediated pulling forces (*esp1 dyn1* cells); (ii) impair MEN activity by inhibiting Cdc15 or Cdc5, two of its essential components (*esp1 cdc15* and *esp1 cdc5* cells); and (iii) inactivate Cdc14 (*esp1 cdc14* cells). In agreement with our hypothesis, most of the *esp1 dyn1*, *esp1 cdc15*, *esp1 cdc5,* and *esp1 cdc14* cells retained a short bipolar spindle configuration for the entire duration of the experiment (**supplementary figure 5**), allowing us to analyze the aMTs of *esp1* mutant cells at their terminal phenotype. aMTs of all the combinations of *esp1* mutants analyzed were significantly longer and more numerous than aMTs of *cdc20* cells, and they resembled the aMTs of the double mutant cells (with an average length of 4µm, 4.2µm, 4µm, 4.5µm, and 4.2µm, and an average number of 3.8 aMT/cell, 4.4 aMT/cell, 4.1 aMT/cell, 4.7 aMT/cell, and 4.3 aMT/cell in *esp1 cdc15*, *esp1 cdc5*, *esp1 cdc14 esp1, cdc5 cdc14,* and *esp1 dyn1* cells, respectively; compared to *cdc20* and *cdc14 cdc5* cells that showed an average aMT length of 2.6µm and 4µm, and an average number of 3.2 aMT/cell and 4.3 aMT/cell, respectively; **figure 3A** and **3B**). Altogether, these data indicate that the activation of the APC/C^Cdc20^ is the molecular event required to promote, directly or indirectly, aMT stabilization at anaphase onset.

**Figure 3.**
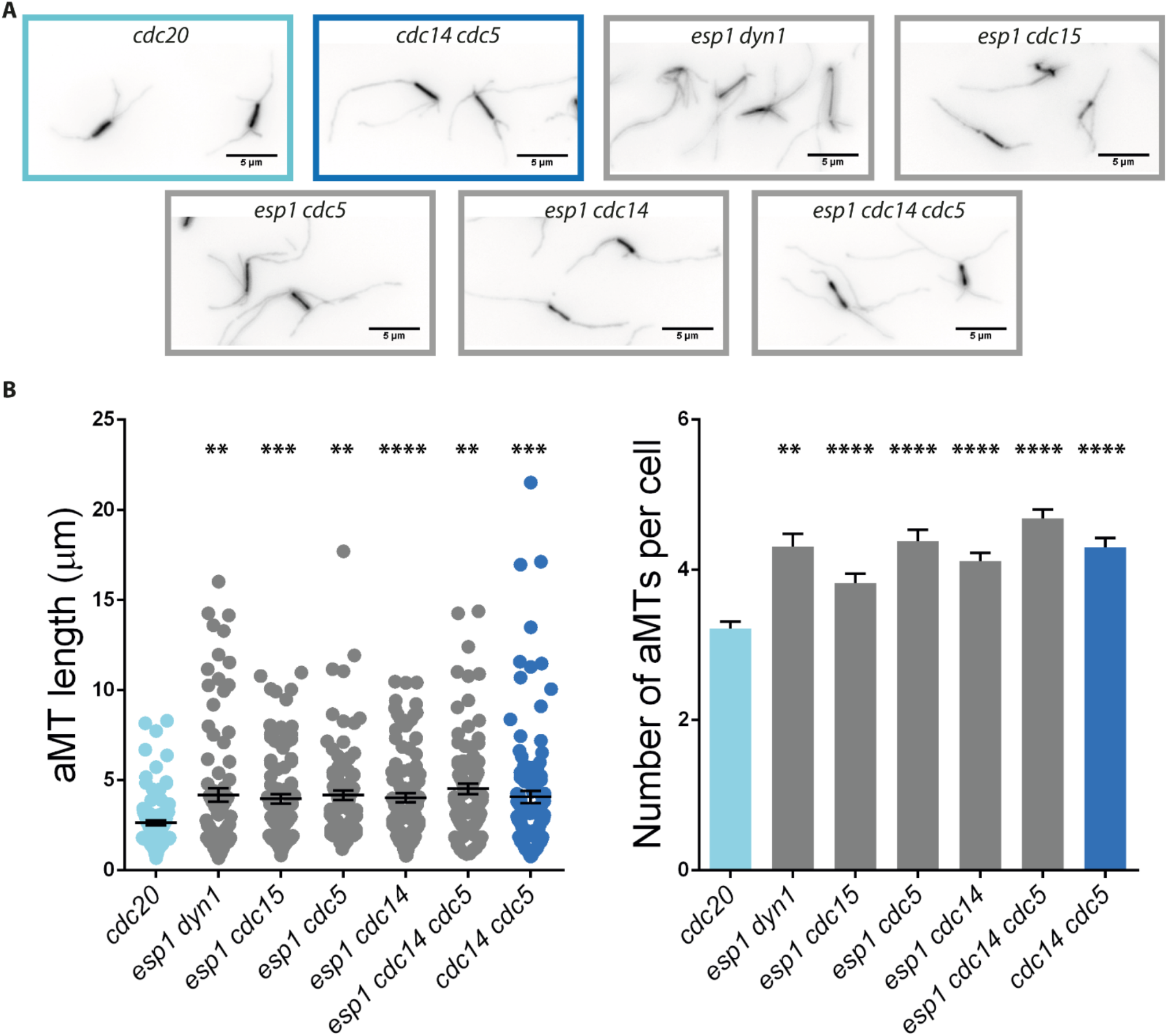
APC/C^Cdc20^ activation is critical for anaphase aMT stabilization. (A-B) *esp-1* (Ry9490), *esp1-1 dyn1*Δ (Ry9516), *esp1-1 cdc15-as1* (Ry9512), *esp1-1 cdc5-as1* (Ry9134*) esp1-1 cdc14-1* (Ry9131), *esp1-1 cdc14-1 cdc5-as1* (Ry9128), and *cdc14-1 cdc5-as1* (Ry1602) cells were synchronized in G1 with α-factor (5µg/ml) and released at 37°C, to inactivate both the *esp1-1* and *cdc14-1* alleles, into fresh YEPD media lacking pheromone and supplemented with 1NM-PP1 analogue 9 (5µM) and CMK (5µM) to inactivate the *cdc15-as1* and *cdc5-as1* alleles, respectively. Cells were scored at their terminal arrest (∼3 hours after the release). (A) Representative images for each mutant at its terminal arrest are shown (scale bar = 5µm). (B) The histograms show aMT length and number of the indicated mutants (n=100; **=p<0.01; ***=p<0.001; ****=p<0.0001; * asterisks denote significant differences according to ordinary One-Way ANOVA and Tukey’s multiple comparisons test).

### The APC/C^Cdc20^ stabilizes aMTs through the degradation of the mitotic cyclin **Clb4**

Given that the most acknowledged function of the APC/C^Cdc20^ complex is to target proteins for proteasomal degradation [42], we searched among its substrates for the one(s) whose degradation has an impact on aMT dynamics. In budding yeast, APC/C^Cdc20^ removes securin Pds1, the critical anaphase inhibitor, at the metaphase to anaphase transition [43] and this is sufficient to trigger spindle elongation and chromosome segregation [43], [44]. To assess whether Pds1 removal is sufficient also for aMT stabilization, we analyzed the impact of Pds1 deletion on metaphase arrested cells. To maintain the short bipolar spindle morphology, which is characteristic of both metaphase and *cdc14 cdc5* cells, we prevented *pMET-CDC20 pds1* cells from elongating their spindle by inactivating Cdc14 and Cdc5 in this mutant background (*pMET-CDC20 pds1 cdc14 cdc5* cells). Simultaneous inactivation of the two proteins does not affect aMT dynamics in metaphase (**supplementary figure 3D** and **3E**). *pMET-CDC20 pds1 cdc14 cdc5* cells were then compared with *pMET-CDC20 cdc14 cdc5* and *cdc14 cdc5 pds1* cells. We found that aMTs of *pMET-CDC20 pds1 cdc14 cdc5* cells are similar in length and number to aMTs of *pMET-CDC20 cdc14 cdc5* cells, but are significantly less and shorter than aMTs of *cdc14 cdc5 pds1* cells (with an average length of 3.7µm, 2.5µm and 2.6µm, and an average number of 3.3 aMT/cell, 2.4 aMT/cell and 2.5 aMT/cell in *cdc14 cdc5 pds1*, *pMET-CDC20 pds1 cdc14 cdc5* and *pMET-CDC20 cdc14 cdc5,* respectively; **supplementary figure 6A**). This result indicates that Pds1 removal *per se* is not sufficient to promote aMT stabilization and points toward other putative candidates.

In higher eukaryotes, in addition to Pds1 removal, the degradation of cyclins is also essential to trigger anaphase entry [45]. In budding yeast, the key cyclin that is targeted for degradation by the APC/C^Cdc20^ is the S-phase cyclin Clb5 [46] and, as such, we wondered whether Clb5 might be the APC/C^Cdc20^ substrate involved in aMT regulation. A role for Clb5 in the control of aMT dynamics is supported by the findings that link the S-phase cyclin to the maturation of the outer plaque of the two Spindle Pole Bodies (SPBs) – the yeast counterpart of centrosomes -, a process that defines their aMT nucleation capacity and, hence, controls aMT morphology [47], [48]. To assess whether Clb5 removal is sufficient to stabilize aMTs, we deleted *CLB5* in a *cdc20* mutant background and probed aMTs at the terminal arrest. We found that the aMTs of *cdc20 clb5* metaphase arrested cells slightly increased both in length and number compared to aMTs of *cdc20* mutant cells (from an average length of 2.4µm and a number of 3 aMT/cell in *cdc20* cells to 3.3µm and 3.4 aMT/cell in *cdc20 clb5* cells, respectively; **supplementary figure 6B**), suggesting that Clb5 removal could be at least partially involved in APC/C^Cdc20^-mediated stabilization of aMTs. A caveat with this analysis is that cells lacking one of the cyclin subunits can undergo compensatory mechanisms. To establish whether this caveat could explain the partial phenotype observed, we assessed the contribution of the S-phase cyclin using the conditional *CLB5-AID* allele. *cdc20-1* and *cdc20-1 CLB5-AID* cells were released from a G1 arrest and when the majority of cells reached metaphase, Clb5 degradation was induced adding Auxin to the media. Also in this case, both the length and number of aMTs only slightly increased in *cdc20 clb5-AID* cells (**supplementary figure 6B**). Although we cannot formally exclude that, in this case, some residual Clb5 activity may account for the intermediate aMT phenotype observed, the convergence of the results obtained with independent *clb5* mutant alleles indicates that Clb5 degradation alone is not sufficient to explain the stabilization of aMTs observed at anaphase onset and a more complex picture has to be envisioned.

Beside degrading the S-phase cyclin Clb5, the APC/C^Cdc20^ initiates the degradation of the main M-phase cyclin Clb2 [45], and possibly all mitotic cyclins, namely Clb1, Clb3 and Clb4. Although, it was reported that Clb2 levels remain high in *cdc14 cdc5* cells [14], as assessed by western blot analysis and consistent with Clb2 degradation requiring Cdc5 activity [49], our mass spectrometric finding that Clb2 and Clb3 levels decrease at the metaphase to anaphase transition prompted us to assess whether other mitotic cyclins contribute to aMT regulation. To this aim and to avoid issues of synthetic lethality (e.g. most *cdc20* mutant alleles are synthetically lethal with *CLB2* deletion (Ref [50] and **supplementary figure 6C**), we tested the consequences of deleting individual cyclin subunits and probed aMTs dynamics in cells arrested with Hydroxyurea (HU). HU reduces the dNTP cellular pool below the levels necessary for DNA replication [51], [52]. This leads to cells arresting in early S-phase or eventually reaching a checkpoint-mediated metaphase arrest, with short bipolar spindles and undivided nuclei. We found that of all tested cyclin subunits, only deletion of Clb4 significantly impacted aMT length and number (**figure 4A**), pinpointing Clb4 as the critical substrate of the APC/C^Cdc20^ affecting aMT stabilization. To confirm this result, we deleted *CLB4* in a *cdc20* background and analyzed aMTs of *cdc20 clb4* cells upon reaching the metaphase arrest (**figure 4B** and **4C**). We confirmed that lack of Clb4 led to abnormally stable metaphase aMTs. aMTs of *cdc20 clb4* cells were as stable as the ones of *esp1 cdc15* and *cdc5 cdc14* double mutant cells (*cdc20 clb4* cells exhibited an average aMT length of 3.9µm and 4.5 aMT/cell, compared to 2.4µm and 3.5µm and 3.2 aMT/cell and 4.4 aMT/cell in *cdc20* and *esp1 cdc15* cells, respectively, **figure 4B** and **4C**). As Clb4 was never formally ascertained as an APC/C^Cdc20^ substrate, to assess the contribution of the APC/C^Cdc20^ in modulating Clb4 kinetics we probed Clb4 levels in S-phase arrested cells following the overexpression of Cdc20 via the galactose-induced promoter (*GAL-CDC20*)[6]. In agreement with what previously reported for APC/C^Cdc20^ substrates (e.g Pds1 [6]), we found that high levels of Cdc20 lead to fast Clb4 degradation (**figure 4D**), on the contrary Clb2, whose degradation is mainly dependent on the APC/C^Cdh1^ complex[6], [53] remains stable and eventually accumulates (**figure 4D**). The findings that: (i) aMTs of *cdc20 clb4* cells phenocopy the ones of *esp1 cdc15* and *cdc14 cdc5* cells, (ii) this phenotype is recapitulated only by the removal of Clb4, and finally (iii) Clb4 is an APC/C^Cdc20^ substrate, confirms Clb4 as the critical APC/C^Cdc20^ substrate whose degradation specifically triggers anaphase-aMTs stabilization.

**Figure 4.**
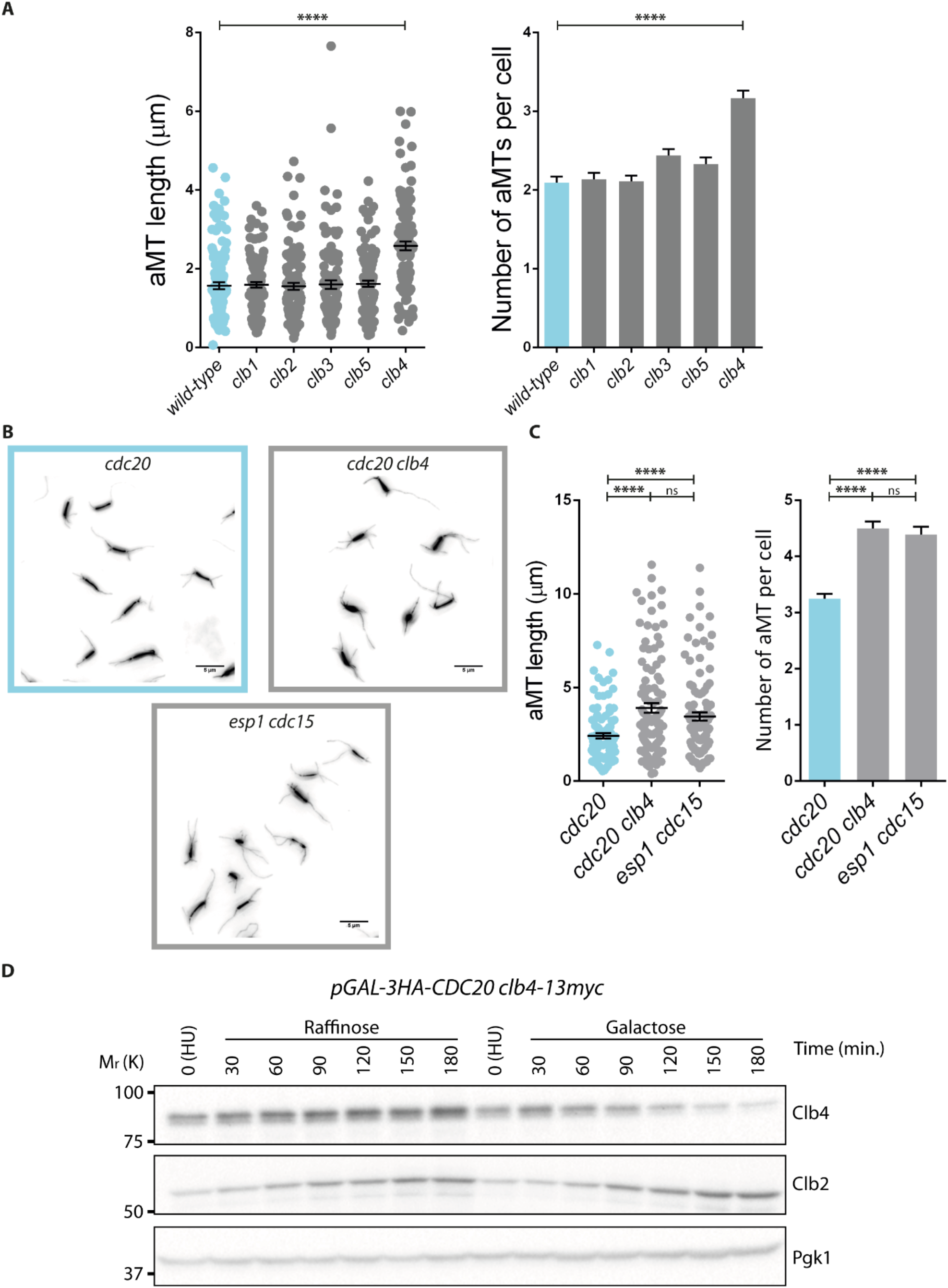
Systematic analysis of putative APC/C^Cdc20^ involved in anaphase aMT stabilization unveils the key role of B-Cyclin Clb4. (A) *Wildtype* (Ry1), *clb1*Δ (Ry5976), *clb2*Δ (Ry20), *clb3*Δ (Ry10741), *clb4*Δ (Ry10824) and *clb5*Δ (Ry445) were arrested in S-phase with HU (10 mg/ml) and analyzed 90 minutes upon reaching the arrest. The graphs show aMT length and number of the indicated genotypes (n=100; ****=p<0.0001; * asterisks denote significant differences according to ordinary One-Way ANOVA and Tukey’s multiple comparisons test). (B-C) *cdc20-AID* (Ry7873)*, cdc20-AID clb4*Δ (Ry10741) and *esp1-1 cdc15-as1* (Ry9512) cells were arrested in G1, released into fresh YEPD media at 37°C to inactivate the *esp1-1* allele, and supplemented with Auxin (500µM) and 1-NM-PP1 analogue 9 (5µM) to inactivate the *cdc20-aid* and *cdc15-as*, alleles, respectively. Cells were analyzed at their terminal arrest (∼3 hours from the release). Representative images (scale bar = 5µm) (B) and graphs for aMT length and number (n=100; *=p<0.05; **=p<0.01; ***=p<0.001; ****=p<0.0001; * asterisks denote significant differences according to ordinary One-Way ANOVA and Tukey’s multiple comparisons test) (C) are shown. (D) *pGAL-3HA-CDC20 clb4-13myc* (Ry10889) cells were arrested in S-phase with HU (10mg/ml) in YEP media supplemented with Raffinose. Upon reaching the arrest (around 180 minutes from HU addition), the culture was split in two. One half was maintained in the same conditions, whereas 2% galactose was added to the other half to induce the expression of Cdc20. Samples were taken at the indicated times. Note: Pgk1 protein was used as an internal loading control in immunoblots. Size markers on the sides of the gel blots indicate relative molecular mass.

To exclude the contribution of other APC/C^Cdc20^ targets, besides cyclins, we also tested: (i) the kinesin motor protein Kip1 [54]; (ii) the Haspin-like kinase Alk2 [55]; (iii) the APC/C^Cdh1^ inhibitor Acm1 [56]; (iv) and the DNA replication-promoting kinase Dbf4 [57]. Individual proteins were depleted in a *cdc20* mutant background and the double mutants were next assessed for aMT morphology. Since, Kip1, Alk2, and Acm1 are non-essential proteins, we probed the consequences of their inactivation using deletion alleles for their encoding genes. Instead, for the essential kinase Dbf4, we employed the thermo-sensitive *dbf4-1* allele. Moreover, to exclude possible compensatory mechanisms mediated by the Alk2-paralog Alk1, we also analyzed *cdc20 alk1 alk2* cells. We found that *cdc20 kip1* and *cdc20 acm1* cells exhibited the partial phenotype already observed in cells lacking Clb5 (a slight increase in aMT length and a similar number of aMTs to their metaphase counterpart; **supplementary figure 6D**), while all the other mutant combinations retained unstable aMTs comparable to aMTs of *cdc20* cells (**supplementary figure 6D**). These findings suggest that, while the main target of the APC/C^Cdc20^ involved in anaphase aMT stabilization is the mitotic cyclin Clb4, a minor role may be played by Clb5, Kip1 and Acm1.

### Multiplexed quantitative phosphoproteomics associates aMT stabilization to de-phosphorylation

To gain insights into the complex molecular mechanism underlying aMT dynamics and guided by the central role played by mitotic cyclin Clb4 in the process, we adopted an unbiased approach and interrogated the proteome and phosphoproteomes of *cdc20*, *cdc14 cdc5* and *cdc15* cells representing metaphase, mini-anaphase and anaphase, respectively.

For the analysis, cells carrying the genotype of interest were arrested in G1 and synchronously released into restrictive conditions. Samples were collected at the terminal phenotype and subjected to tandem mass tags-based multiplexed quantitative phosphoproteomics. Our dataset covered 4,641 proteins (**supplementary table 2**) and identified 5,324 phospho-sites (**supplementary table 3**). To establish that the interrogated mutants are *bona fide* representatives of metaphase, early and late anaphase, hence provide us with a snapshot of the proteome and phosphoproteome of these cell cycle phases, we initially looked at CDK1 substrates (Fin1 [58], Ase1 [10] and Orc6 [58]) whose de-phosphorylation status is known to change in this cell cycle window. We identified 2 out of 5 putative CDK1 residues reported for Fin1 [59] (S36, T68; **supplementary figure 7A**), 4 out of 7 for Ase1 [10] (T55, S64, S198, S803; **supplementary figure 7B**) and 3 out of 4 for Orc6 [60] (S106, S116, T146; **supplementary figure 7C**). Consistent with Fin1 being de-phosphorylated in early anaphase, Ase1 in mid anaphase and Orc6 at mitotic exit, when we compared the *cdc20* and *cdc15* datasets, we found that all Fin1 residues, 2 out of 4 Ase1 residues and no Orc6 residues were de-phosphorylated (**supplementary figure 7**). Moreover, when the *cdc14 cdc5* dataset was included in the analysis and compared to the other two datasets individually, also residues whose de-phosphorylation is Cdc14-dependent were identified (i.e., Ase1^S803^; **supplementary figure 7**).

Having established that our mutants are a good proxy for the mitotic cell cycle phases of interest, we assessed global phosphorylation changes. We identified 974 phosphosites that changed their status when going from metaphase to anaphase, as assessed by comparing *cdc20* with *cdc15* cells (**figure 5A**). Among the 974 phosphosites, 197 were phosphorylated (69 of which reside in the minimal CDK1 consensus site) and 777 were de-phosphorylated (of which 407 are putative CDK substrates), showing an overall trend in favor of de-phosphorylation. Since aMT regulation is characteristic of anaphase and independent of Cdc14 and Cdc5, we restricted the analysis by comparing the phosphoproteome of *cdc20* cells with the combination of the datasets of *cdc14 cdc5* and *cdc15* cells (**figure 5B**). By these means, we identified 340 phosphosites that changed from metaphase to anaphase, of which 81 turned out to be phosphorylated (34 of which reside in the minimal CDK1 consensus site) and 259 were de-phosphorylated (of which 160 are putative CDK substrates). The changes of these phosphorylation sites were unlikely caused by changes in protein abundance as the respective protein amounts remained largely unchanged (**figure 5A** and **5B**). Our data, in contrast with reported finding that in anaphase the overall number of phosphorylated and de-phosphorylated residues is similar [13], evidence a bias towards de-phosphorylation for anaphase cells. A reason for this discrepancy may be the experimental set up, while in the previous study the analysis was performed in cells undergoing a synchronous cell cycle, here we employed homogenously arrested populations.

**Figure 5.**
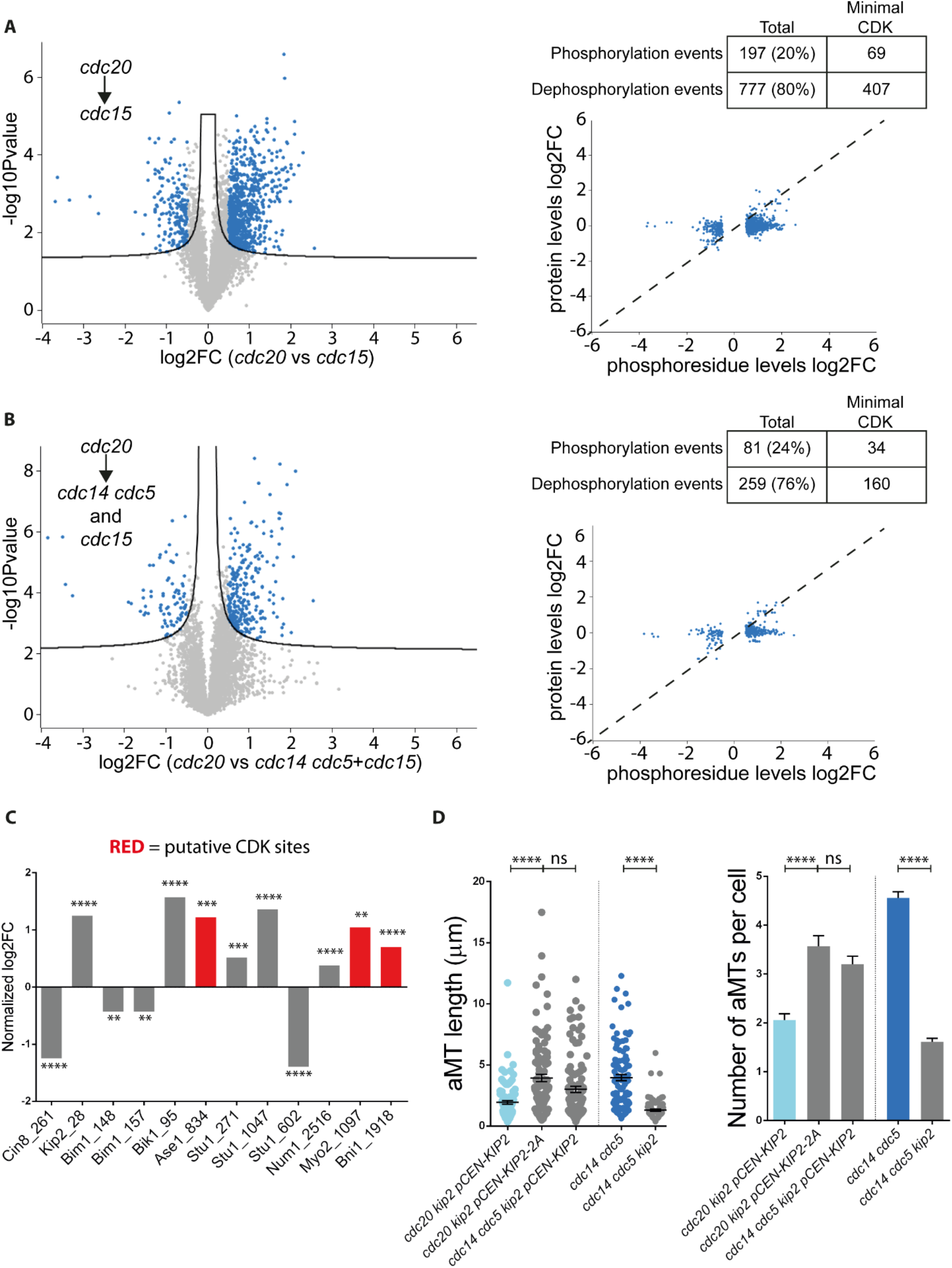
Modeling the metaphase to anaphase transition by phospho-proteomics unveiled numerous residues that may be involved in anaphase aMT stabilization. (A-C) *cdc20-AID* (Ry4853)*, cdc14-1 cdc5-as1* (Ry1602), and *cdc15-as1* (Ry1112) cells were arrested in G1 and released at 37°C, to inactivate the *cdc14-1* allele, into fresh media supplemented with Auxin (500 µM), the 1NM-PP1 analogue 9 (5µM) and the CMK (5µM) inhibitors to inactivate the *cdc20-AID*, *cdc15-as1*, and the *cdc5-as1* alleles, respectively. Cells were harvested at their terminal arrest (∼3 hours after the release) to extract proteins and processed for TMT mass spectrometry analysis with three biological replicas for *cdc20-AID* and *cdc14-1 cdc5-as1* cells and two biological replicas for *cdc15-as1* cells. (A) The volcano plot represents the log2 fold change and the respective *–log10 p-value* were calculated comparing the values obtained for each phospho-residue in *cdc20-AID* and *cdc15-as1* cells. To identify the phospho-residues that were significantly different between the two mutant cells, a FDR of 0.001 (S0 at 0.05) was applied, with an additional cut-off of a *log2* fold change <-0.5 or >0.5. The log2 fold changes of the phospho-residues obtained with this analysis were also plotted against the log2 fold changes of the corresponding proteins and scored for those that belong to the minimal CDK consensus (S/TP). (B) For each phospho-residue obtained, the log2 fold change and the respective log10 p-value were calculated comparing *cdc20-AID* with *cdc14-1 cdc5-as1* and *cdc15-as1* cells together. The generated data were processed as in (A). (C) The graph shows the phospho-residues that belong to proteins already linked to the regulation of aMT dynamics and their log2 fold changes as calculated in (B) and normalized by the log2 fold change of the protein to account for protein abundance change. Only residues with a log2 fold-change of at least 0.4 and a significant p-value are shown. The residues that belong to the minimal CDK consensus are shown in red. (**=p<0.01; ***=p<0.001; ****=p<0.0001). (D) *cdc20-AID kip2Δ pCEN-KIP2* (Ry10858), *cdc20-AID kip2Δ pCEN-KIP2S63AT275A* (Ry10861), *cdc14-1 cdc5-as1 kip2Δ pCEN-KIP2* (Ry10857), *cdc14-1 cdc5-as1 kip2*Δ (Ry10844) and *cdc14-1 cdc5-as1* (Ry1602) arrested in G1 and released at 37°C, to inactivate the *cdc14-1* allele, into fresh media supplemented with Auxin (500 µM) and the CMK (5µM) inhibitors to inactivate the *cdc20-AID* and the *cdc5-as1* alleles, respectively. Cells were analyzed at their terminal arrest (∼3 hours after the release) and both aMT length and number was measured. (n=100; ****=p<0.0001; * asterisks denote significant differences according to ordinary One-Way ANOVA and Tukey’s multiple comparisons test).

To identify possible regulators of aMT dynamics, we scrutinized the phosphorylation status of residues within proteins known to be involved in microtubule regulation [31], [61]–[63], namely five kinesin motor proteins (Cin8 [64], Kip1 [64], Kip2 [65], Kip3 [66], and Kar3 [67]), the single minus-end directed Dyn1 protein, and five MAPs (Bim1 [68], Bik1 [69], Ase1 [70], Stu1 [71] and Stu2 [72]). Of note, all proteins are equally abundant in the three mutant strains except for Kip1, which, being an APC/C^Cdc20^ substrate, shows an abundance pattern that is typical for ubiquitin ligase targets. Given that the activity of these proteins is often modulated by microtubule-cortex interaction [62], [73], [74], we extended our analysis to proteins that mediate the connection between aMTs and the cellular cortex, such as the cortical receptor Num1 [75], the actin motor protein Myo2 [33] and the actin nucleator Bni1 [62]. Interestingly, we found that Cin8, Kip2, Bim1, Bik1, Ase1, Stu1, Num1, Myo2, and Bni1 showed at least one residue with a different phosphorylation status between metaphase and anaphase arrested cells (**figure 5C**) with a bias in favor of de-phosphorylation. To validate our analysis we chose the kinesin Kip2. Kip2 has a microtubule polymerization function and it is negatively regulated by phosphorylation [23], [76]. This inhibitory phosphorylation is mediated by the GSK3 kinase and requires priming by an additional kinase, of which CDK is one candidate, at S63 and T275 Kip2 sites [23]. To test if aMTs are kept unstable in metaphase by CDK-mediated phosphorylation of Kip2, we probed aMTs in *cdc20 kip2-S63AT275A* mutant cells (henceforth *kip2*-2A) and found that they pheno-copy the ones of the double mutant (**figure 5D**). Next, we confirmed that anaphase stabilization of aMTs requires Kip2 activity by deleting *KIP2* in *cdc14 cdc5* double mutant cells. As expected we found that lack of the kinesin reduced both aMTs length and number (**figure 5D**). These results identify Kip2 as a Clb4 substrate and pave the way for further testing of putative substrates/sites within proteins of which Ase1 (S834), Myo2 (T1097) and Bni1 (T1918) are an example.

### Dynamic aMTs and Cdc5 guide proper spindle orientation

What is the physiological significance of this regulation? The identification of Clb4 as central to the regulation of aMT dynamics contributes to our understanding of the mechanism by which budding yeast properly aligns its mitotic spindle along the mother-bud axis, an essential requisite for survival. Schiebel and colleagues [17] proposed that Clb4 in complex with CDK1 facilitates spindle alignment by mediating the interactions of aMTs with the bud cortex and enables the turnover of established aMT-bud cortex attachments^3^, and our data suggest that it does so by controlling aMT stability. This model allows clear predictions: (i) stable microtubules should exhibit an increased residence time at the cortex and (ii) premature aMT stabilization should result in an increased amount of wrong attachments.

To assess if the increased aMT stability observed in the double mutant cells reflects changes in the dynamics of aMT-cortex interactions, we analyzed the behavior of aMTs at the cellular cortex in *cdc20* and *cdc14 cdc5* arrested cells carrying a GFP-tagged Tub1 fusion by live-cell imaging, and measured the time that each individual aMT spent contacting the cellular cortex. In agreement with our hypothesis, aMTs of *cdc14 cdc5* cells remain bound to the cellular cortex longer than aMTs of *cdc20* cells (**figure 6A** and **6B**; on average, aMTs of *cdc14 cdc5* and *cdc20* cells remain close to the cortex during 77% and 39% of the analyzed time, respectively), thus indicating that the dynamics of aMT-cortex connections are altered in the double mutant cells.

**Figure 6.**
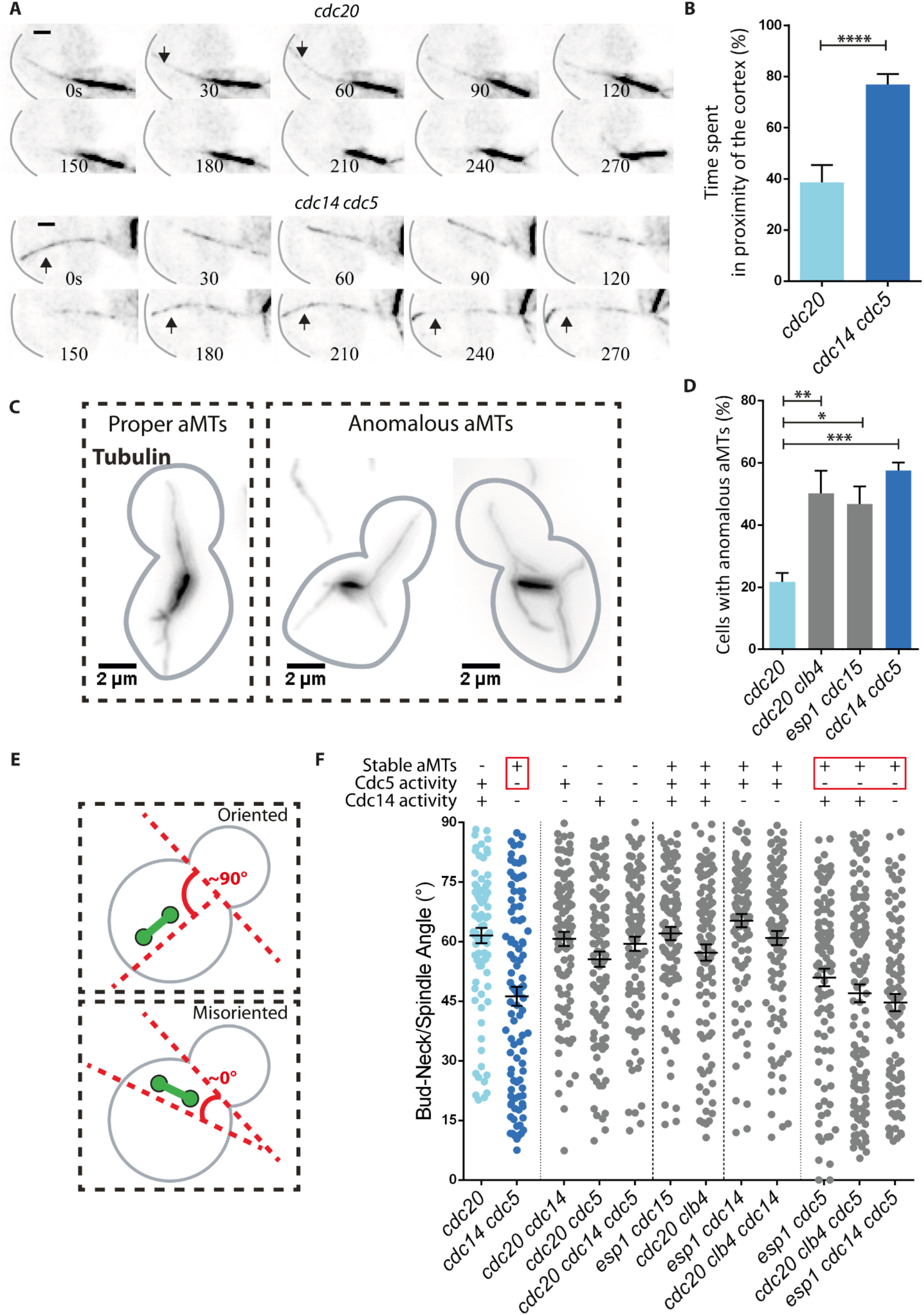
Dynamic aMTs and Cdc5 guide spindle positioning. (A-B) *cdc20-AID* (Ry7732) and *cdc14-1 cdc5-as1* (Ry3256) cells harboring a *TUB1-GFP* fusion were synchronized in G1 and released at 37°C into fresh media supplemented with Auxin (500µM), the 1NM-PP1 analogue 9 (5µM) and the CMK (5µM) inhibitors to inactivate the *cdc20-AID*, *cdc15-as1* and *cdc5-as1* alleles, respectively. Approximately 3 hours after the release, cells were moved to a CellASIC ONIX plate and z-stack time-lapse images of spindle microtubules were acquired every x second per x minute to determine aMT length. (A) Time-lapse images from a representative cell are shown for both *cdc20-aid* and *cdc14-1 cdc5-as1* strains (scale bar = 1µm). Black arrows indicate the frames when the highlighted aMTs were in close proximity with the cellular cortex. (B) The histograms show the quantification of the time that each aMT spent in proximity with the cortex (n= 100; ****=p<0.0001; * asterisks denote significant differences according to ordinary One-Way ANOVA and Tukey’s multiple comparisons test). (C-F) *cdc20-AID* (Ry4853), *cdc14-1 cdc5-as1* (Ry1602), *pMET-CDC20 cdc5-as1* (Ry3209)*, pMET-CDC20 cdc14-1 cdc5-as1* (Ry3201)*, pMET-CDC20 cdc14-1* (Ry3204)*, esp1-1 cdc15-as1* (Ry9512)*, cdc20-AID clb4*Δ (Ry10741)*, esp1-1 cdc14-1* (Ry9131)*, esp1 cdc5-as1* (Ry9134), and *esp1-1 cdc14-1 cdc5-as1* (Ry9128) cells were synchronized in G1 and released at restrictive conditions for each mutant. Cells were analyzed at their terminal arrest (∼3 hours after the release). (C) Representative images of cells with proper or anomalous aMTs, a distinction based on aMT direction, are shown (scale bar = 2µm). (D) The histograms show the quantification of anomalous aMTs in the four strains (n=100; ***=p<0.001; * asterisks denote significant differences according to unpaired t-test). (E) Schematic representation of the angles generated by the bud-neck and the spindle, an indicator of spindle orientation. The closer the angle is to 90°, the more the spindle is properly oriented toward the mother-bud axis; vice versa, the more the angle is close to 0°, the more it is misoriented. (F) The dot plot shows the bud-neck/spindle angles measured in the cells of the indicated genotype. Each data point in the dot plot represents a single cell. Note that an average angle of 45° means a complete randomization of spindle orientation. (n=100 per strain).

To assess if premature aMT stabilization leads to an increased amount of wrong attachments, we probed aMT directionality. Proper cortical attachments foresee that aMTs originating from the bud-directed SPB enter the bud, while the others point to the cortex of the mother cell. If stabilization prevents error correction, then mutants with prematurely stable microtubules should manifest directionality problems, with aMTs originating from two different SPBs pointing to the same cellular compartment, and aMTs originating from a single SPB pointing to both mother and daughter cells (in our analysis, both possibilities were grouped under the category of abnormal aMTs). Indeed, when we scrutinized aMT directions in *cdc20*, *cdc20 clb4 esp1 cdc15* and *cdc14 cdc5* cells, we found that all mutants with stable aMTs (*cdc20 clb4 esp1 cdc15* and *cdc14 cdc5)* showed an increased percentage of abnormal aMTs when compared to *cdc20* mutants (on average, 22% of *cdc20* cells, 50% of *cdc20 clb4* cells, 47% of *esp1 cdc15* cells and 58% of *cdc14 cdc5* cells showed anomalous aMTs; **figures 6C** and **6D**).

Since proper attachment of aMTs with the bud cortex directs the spindle toward the emerging bud and orients it to the polarity axis up to anaphase onset, when – concomitantly to spindle elongation – they pull only one of the two poles across the bud-neck [77]–[79], we asked whether anomalous aMTs affect spindle orientation. Spindle orientation was analyzed by measuring the angle formed by the spindle and the bud-neck in *cdc20*, *cdc14 cdc5*, *esp1 cdc15*, and *cdc20 clb4* cells. The closer the angle is to 90°, the more the spindle is correctly oriented toward the polarity axis, while the closer the angle is to 0°, the more the spindle is misoriented (**figure 6E**). We found that, while most metaphase arrested cells correctly orient their spindle, *cdc14 cdc*5 cells do so in a random manner, as indicated by the mean bud-neck/spindle angle of 45° (**figure 6F**). If this finding supports the idea that stable aMTs lead to improper aMT-bud cortex attachments and, ultimately, spindle positioning defects, then the observation that, in *esp1 cdc15* cells and *cdc20 clb4* spindles were properly oriented (**figure 6F**) indicates that astral microtubule stability predisposes to, but it is not sufficient to cause spindle orientation defects.

Since among all the tested mutants only *cdc14 cdc5* manifested spindle orientation defects, we wondered if Cdc14 and/or Cdc5 inactivation could account for the phenotype. To test this hypothesis, spindle orientation was analyzed in *cdc20 cdc14*, *cdc20 cdc5*, and *cdc20 cdc14 cdc5* cells, and we found that all tested strains carry properly oriented spindles (**figure 6F**), hence indicating that Cdc14 and Cdc5 inactivation it is not sufficient to impair spindle orientation. However, the finding that *esp1 cdc5*, *esp1 cdc14 cdc5* and *cdc20 clb4 cdc5* cells, in which aMTs are stable and Cdc5 is inactivated, showed orientation defects (**figure 6F**), suggests that unstable aMTs and Cdc5 activity are redundantly required for proper spindle orientation.

## Discussion

Despite the importance of astral microtubules in controlling spindle stability and orientation, their regulation remains largely understudied. Here we report that, similarly to kinetochore and interpolar microtubules, aMT dynamics are also intrinsically regulated in a cell cycle dependent manner. More precisely, they remain unstable up to metaphase and are next stabilized as soon as cells enter anaphase. This finding is consistent with their role in searching for cortical anchor sites and correcting erroneous attachments in metaphase and next in stabilizing the binding of the mitotic spindle with the cortex to properly guide spindle positioning and elongation in anaphase.

Central to the regulation of astral microtubule dynamics are the Clb4-CDK1 and APC/C^Cdc20^ complexes – two evolutionary conserved mitotic machineries. More precisely, the presence of Clb4 is sufficient to render aMTs unstable up to metaphase likely by directly or indirectly introducing one or multiple inhibitory phosphorylation events in factors affecting microtubule stability of which Kip2 is an example. *Viceversa*, the enforcement of bindings observed at anaphase and dependent on aMT stabilization is dictated by the APC/C^Cdc20^–mediated degradation of the mitotic cyclin Clb4. Of note Clb4 is unique among all mitotic cyclins in its ability to affect aMT dynamics as indicated by the finding that only its removal is sufficient to stabilize aMTs in cell cycle phases when they are normally unstable. Finally, aMTs stability in anaphase is counteracted by a mechanism that likely integrates two anaphase specific requirements, namely spindle elongation and stable aMT-cortex links.

The identification of a kinase as central to aMT regulation, immediately calls for the phosphatase required to reverse these Clb4-CDK1-mediated phosphorylation events. If in yeast the main mitotic CDK-counteracting phosphatase is Cdc14, our data indicate that at least one additional phosphatase may be involved. A possible candidate is Glc7 (the sole PP1 catalytic subunit in yeast) in combination with its regulatory subunit Bud14. Indeed, overexpression of Bud14 increases aMT length [80] in a way reminiscent of cells lacking Clb4. The involvement of the Glc7-Bud14 complex is also supported by its localization. Both Glc7 and Bud14 accumulate at bud cellular cortex [80], [81], an ideal place to counteract the activity of the Clb4-CDK1 complex, which mainly localizes at the plus-end of aMTs directed toward the bud [82]. Importantly, this interplay between Clb4-CDK1 and Bud14-Glc7 at the interface between aMT plus-ends and the bud cellular cortex, besides satisfying the requisite of the yet-to-identify “spatial cue” - necessary for the specific stabilization of bud-directed aMTs - reported by Barral and collegues [83], it also integrates intrinsic and external signals.

The observation that aMT dynamics are under the control of CDK1 and APC/C^Cdc20^ activities makes it likely that the molecular mechanism worked out in yeast is retained in vertebrates. At least three pieces of evidence support this hypothesis. First, in all vertebrates CDK1-CyclinB activity is high in metaphase and decreases at anaphase onset due to APC/C^Cdc20^-mediated degradation of the cyclin B subunit [45]. Evidences exist that in human cells, CDK1-CyclinB activity destabilizes aMTs in prometaphase by phosphorylating the EB1-dependent microtubule plus-end tracking protein GTSE1 [22]. Second, the knowledge that Dyn1 activity in mammalian cells is low in metaphase and high in anaphase [18], [21], [84], and, at least in budding yeast, this increase in activity is directed by aMT stabilization [32] supports the possibility that aMT are stabilized in anaphase. Third, aMTs are often reported as shrinking during human cell anaphase, thus suggesting that a mechanism counteracting aMT stability connected to spindle elongation/cortex proximity exists also in vertebrates.

Moreover, having identified Cdc5 as a novel regulator of spindle positioning redundant to aMT, provides an explanation as to why this class of microtubules it is finely regulated also in yeast, where they have been considered non-essential based on the knowledge that they are dispensable for spindle elongation. Interestingly, Plk1, the human homologue of Cdc5, has also been intensively linked to the regulation of spindle positioning [85]. Indeed, Plk1-dependent phosphorylation of several components of the cortex platform that bind to aMTs is required to correctly orient and position the spindle along the cleavage plane [86]. Investigating how Cdc5 influences these processes may further clarify the role of the kinase in higher eukaryotes and shed light into how it coordinates spindle positioning with the plethora of mitotic events, under its control. Taken together our work legitimates yeast as an optimal model system to investigate astral microtubule regulation and spindle positioning at molecular level. Besides the fact that it shows only few aMTs (1-6 aMTs per cells) [77], easily distinguishable from other types of microtubules and singularly traceable over time [30], our mass spectrometric analysis clearly illustrates the advantages of exploiting the power of yeast genetics to clarify complex molecular processes.

An important corollary of our study is uncovering the APC/C^Cdc20^ complex as the master “*choreographer*” of late mitotic events. Besides determining the point of no return for mitotic exit by controlling cohesin removal, the APC/C^Cdc20^ precisely coordinates, in time and space, the sequence of events required for the faithful execution of mitosis. It does so by sequentially impacting on the stability of the three classes of spindle microtubule - astral, kinetochore and interpolar - each of which it is intimately linked to a key cell cycle process, namely spindle positioning, cohesin cleavage and chromosome segregation. More precisely, active APC/C^Cdc20^ targets several substrates for degradation, including Clb4, Pds1 and other mitotic cyclins (**figure 7**; Step 1). Degradation of Clb4 is the signal for aMT stabilization (**figure 7**; Step2). Pds1 degradation unleashes the protease Esp1 (**figure 7**; Step 2) – the yeast homologue of separase. Esp1 has a dual role, on the one hands it cleaves the cohesin subunit Scc1 [40], [87], thereby prompting cohesin removal, which in turns affects kMT dynamics, by removing the opposite forces to spindle pulling [88] (**figure 7**; Step 3). On the other hands it promotes, as a component of the FEAR network [37], the transient activation of Cdc14 (**figure 7**; Step 3). Cdc14, next acts on motors and MAPs to stabilize iMTs [8], [14], [89], thereby promoting spindle elongation (**figure 7;** Step 4). The time delay imposed by the incremental number of molecular events impacting specific classes of spindle microtubule (**figure 7**) guarantees that proper spindle positioning precedes sister chromatid separation (cohesin cleavage) which in turns precedes their segregation (spindle elongation). Can this logic of mitotic progression stand true in vertebrates? We already mentioned data supporting a regulation of astral microtubules mediated by CDK1 activity [22]. In respect to chromosomes movement toward the poles (anaphase A) evidence exists that kMT de-polymerization is coupled to cohesin cleavage [90]. Finally spindle elongation, hence chromosomes movement to the opposite poles, relies on the assembly of the central spindle – the Metazoan equivalent of the yeast spindle midzone and this process requires, as in yeast, interpolar microtubule stabilization and bundling. The latter is promoted by the centralspindlin complex - comprising the *Caenorhabditis elegans* ZEN-4 (mammalian orthologue MKLP1) kinesin-like protein and the Rho family GAP CYK-4 (MgcRacGAP) – which in turn is regulated by phosphorylation and de-phosphorylation events of the kinesin motor domain [91]. It has been reported that this phospho-regulation is mediated, directly or indirectly, by a proline-directed kinase-phosphatase switch [91] with Cdc14 being a likely candidate. Indeed, the two human homologues of Cdc14 bind to microtubules and promote both their stabilization and bundling activity during mitosis [92]. Based on the knowledge that in human cells a CDK counteracting phosphatase(s), other than Cdc14 homologues, whose activation necessitates proteasomal activity [93], drives mitotic exit, makes it likely that the stepwise regulation of late mitotic events dictated by the APC/C^Cdc20^ is an evolutionary conserved mechanism of regulation.

**Figure 7.**
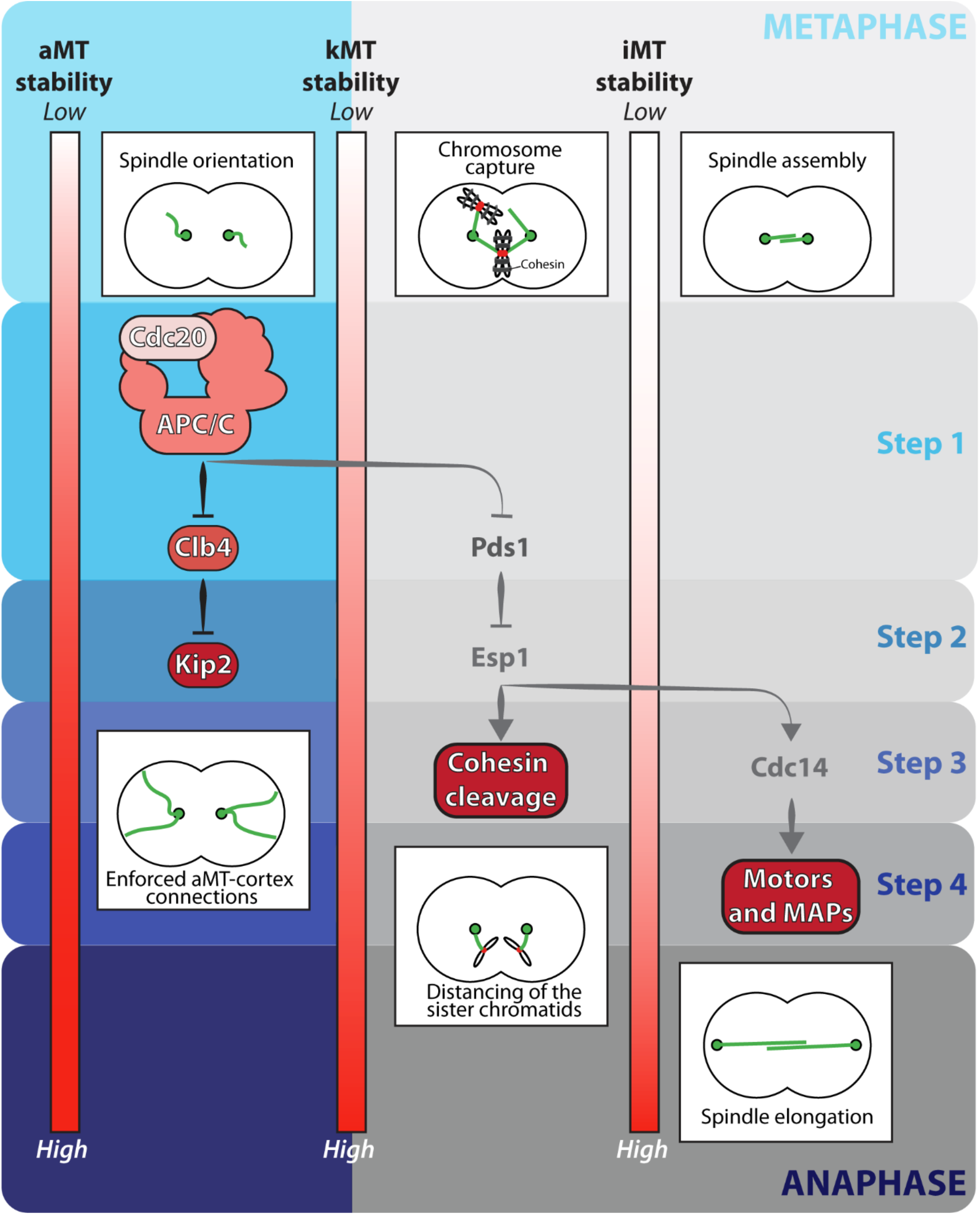
A model for the timely regulation of late mitotic events by the APC/C^Cdc20^. Proper rearrangement of spindle microtubule dynamics coordinates sister chromatid segregation with other late mitotic events. The APC/C^Cdc20^ directs this process by triggering a signaling cascade in which individual steps lead to the regulation of a single class of spindle microtubules. Step 1: The APC/C^Cdc20^ targets for degradation different substrates, including the mitotic cyclin Clb4 and the securin Pds1. Step 2: Clb4 degradation by de-phosphorylating – directly or indirectly-the kinesin-like protein Kip2 promotes aMT stabilization. Instead, Pds1 degradation unleashes the separase Esp1. Step 3: Esp1 cleaves the cohesin subunit Scc1 and activates the Cdk-counteracting phosphatase Cdc14, meanwhile kMTs retract to the poles. Step 4: Cdc14-mediated de-phosphorylation of different motors and MAPs triggers stabilization of iMTs.

## Materials and Methods

### Yeast strains

All strains used in this study are isogenic to W303 and are listed in **supplementary table 1**.

### Growth conditions

Cell cycle arrest and synchronization experiments were performed as previously described[94]. Yeast cells were grown in yeast extract peptone (YEP) media or indicated synthetic medium. Carbon sources (glucose, raffinose or galactose) were used at a final concentration of 2%. Cells were arrested in G1 phase with 5µg/ml α-factor (Genscript) or in S phase with 10 mg/ml HU (Sigma). To release cells from the induced arrest, cells were washed with 5 volumes of fresh medium and transferred into medium lacking the drug. Both the *cdc5-as1* inhibitor CMK and the *cdc15-as1* inhibitor 1NM-PP1 analogue 9 were used at a final concentration of 5µM. Auxin was used at a final concentration of 500 µM.

### Immunostaining

Indirect in situ immunofluorescence methods and antibody concentrations were performed as previously described[24]. Spindle microtubules were detected by α-tubulin immunostaining with the YOL34 monoclonal antibody (MCA78G, AbD Serotec, used at 1:100 dilution) followed by indirect immunofluorescence using FITC-conjugated donkey anti-rat antibody (712-095-153, Jackson ImmunoResearch Laboratories, used at 1:100 dilution).

### Astral microtubule analysis in fixed cells

Images of stained cells were acquired with an upright LEICA DM6 B microscope with a 100X/1.40 oil UPlanSApo ∞/0.17/DFN 25 Olympus objective and Andor Zyla.4.2P camera using Leica Application Suite X software. Optimized z-stacks were taken to cover a thickness of 6.1µm. Image analysis was performed using the “Fiji Is Just ImageJ” (FIJI) software. aMT length was measured in three dimensions using the FIJI plug-in “simple neurite tracer”. To properly score the aMT number, the presence of abnormal aMTs and the bud-neck/spindle angles, Z-series were collapsed into a maximum-intensity two-dimensional projection using the Z-project function. The bud-neck was defined based on the DIC image of the cell.

### Astral microtubule analysis in live cells

50µl of arrested cells carrying a GFP-tagged Tub1 fusion were collected at OD_600_ = 0.2 - 0.4 and loaded in a CellASIC ONIX plate for haploid yeast cells (Millipore). This procedure allows a constant addition of fresh medium to the culture and prevents cellular movements during image acquisition. Images were acquired every 10 seconds for a total of 5 minutes with Nikon Eclipse Ti inverted microscope with a 100X/1.40 oil Olympus objective and Andor Zyla sCMOS camera using the NIS software version 5.10.00. At each time point, 17 z-stack images were taken (0.4µm apart) covering a total thickness of 6.8µm. Image acquisition was followed by a deconvolution process automatically performed by the Huygens software. Image analysis was performed using FIJI. To reduce the complexity and the noise of the analysis, aMT length measurements were performed using the FIJI plug-in “simple neurite tracer” in maximum-intensity two-dimensional projection using the Z-project function. As described in ref.[30], different events were identified: (i) polymerization events - defined as an increase in microtubule length by at least 0.5µm across a minimum of three time points; (ii) depolymerization events - defined as a decrease in microtubule length by at least 0.5 µm across a minimum of three time points; and (iii) pause events - defined as net changes in microtubule length less than 0.5µm across a minimum of 3 time points. Next, catastrophe and rescue events were calculated as previously described[30], [31]: catastrophe frequencies were calculated by dividing the number of polymerization-to-depolymerization transitions by the total time of all growth events; rescue frequencies were calculated by dividing the number of depolymerization-to-polymerization transitions by the total time of all shrinkage events. aMT-cortex connection was evaluated based on the DIC image of the cell.

### Immunoblot analysis

Cells were lysed in 50mM Tris pH 7.5, 1mM EDTA, 1mM *p*-nitrophenyl phosphate, 50mM dithiothreitol, 1mM phenylmethylsulphonyl fluoride, and 2 µgml^−1^ pepstatin with glass beads and boiled in 1x sample buffer. The primary antibodies used were 9E10 monoclonal anti-myc (CVMMS-150R-1000, Covance, used at 1:1,000 dilution) to detect Clb4-13myc; anti-Clb2 (Y-180; SC-9071, Santa Cruz Biotechnology, used at 1:1,000 dilution) to detect Clb2; anti-Pgk1 (A-6457, Molecular Probes, used at 1:5,000 dilution) to detect Pgk1; Goat anti-rabbit IgG (H + L)– HRP conjugate (170-6515, Bio-Rad, used at 1:10,000 dilution) and goat anti-mouse IgG (H + L)–HRP conjugate (170-6516, Bio-Rad, used at 1:10,000 dilution) were used as secondary antibodies to visualize proteins using chemiluminescence (ECL, GE Healthcare).

### Statistical analysis

Depending on the experiment and as indicated in the figure legends, P values were determined by unpaired Student’s t-test or One-Way Anova - Tukey’s multiple comparisons test using the GraphPad Software. P value of less than 0.05 was considered statistically significant (∗ = P < 0.05; ∗∗ = P < 0.01; ∗∗∗ = P < 0.001; ∗∗∗∗ = P < 0.0001). In graphs, averages ± S.E.M. (Standard Error of the Mean) is normally shown.

### Repeatability of the experiments

Each experiment in the manuscript has been repeated at least three times. The results were highly reproducible.

### LC-MS/MS analysis

Sample preparation followed a previously reported procedure [95]. Digested samples were labeled with TMT-11plex reagents (ThermoFisher, 90406, A34807) in the following order: 126 (*cdc15-as1*), 127n (*cdc15-as1*), 129n (*cdc20-AID*), 129c (*cdc20-AID*), 130n (*cdc20-AID*), 130c (*cdc14-1 cdc5-as1*), 131 (*cdc14-1 cdc5-as1*), and 131c (*cdc14-1 cdc5-as1*). Proteomic and phosphoproteomic data were collected on an Orbitrap Fusion and an Orbitrap Lumos mass spectrometer (ThermoFisher Scientific), respectively. An on-line real-time search algorithm (Orbiter) was used in proteomic data collection as described previously[96], [97]. MS raw files were initially converted to mzXML and monoisotopic peaks were re-assigned using Monocle [98]. Database searching with SEQUEST included all entries from the Saccharomyces Genome Database (SGD, 2014). Peptide-spectrum matches (PSMs) were adjusted to 1% false discovery rate (FDR)[99]. Protein-level FDR was filtered to the target 1% FDR level. Phosphorylation site localization was determined using AScore algorithm [100] and filtered at 13 (95% confidence). For TMT reporter ion quantification, a 0.003 Da window around the theoretical m/z of each reporter ion was scanned, and the most intense m/z was used. Reporter ion intensities were adjusted to correct for the isotopic impurities of the TMT reagents according to the manufacturer’s specifications. Peptides were filtered for a summed signal-to-noise of 100 across all channels. For each protein, peptide TMT values were summed to create protein quantifications. To compensate for differential protein loading within a TMT plex, the summed protein quantities were adjusted to be equal within the plex. Phosphorylation site quantifications were also normalized by correction factors generated in this process to account for protein loading variance. For each protein or phosphorylation site within a TMT plex, the signal-to-noise value was scaled to sum to 100 for subsequent analysis. Unpaired student’s t test was performed in Perseus (version 1.6.15.0) to identify the phospho-residues that were significantly different between the different mutant cells. An FDR of 0.001 (S0 at 0.05) was applied, with an additional cut-off of a log2 fold change <-0.5 or >0.5. To identify the phospho-residues likely phosphorylated by CDK, and the phospho-peptides were filtered using the minimal CDK consensus (S/TP).

## Acknowledgments

In memSackory of Angelika Amon an unmatchable mentor and dear friend. We thank Angelika Amon, Dana Branzei, Marco Muzi-Falcone and Orna Cohen-Fix for strains and reagents; Wessen Maruwge for English language editing; Elmar Schiebel, Marina Mapelli, Ian Cheeseman and members of the R.V. laboratory for critical discussions and for critical reading of the manuscript.

## Author contributions

F.Z. and R.V. designed the research project; F.Z., and J.L. performed the research; C.V. provided support to the experimental work; R.V and S.P.G. provided resources; F.Z. and J.L analyzed the data; F.Z. and R.V. wrote the paper. All authors have read and approved of the manuscript.

## Supplementary Materials

**Fig. S1.**
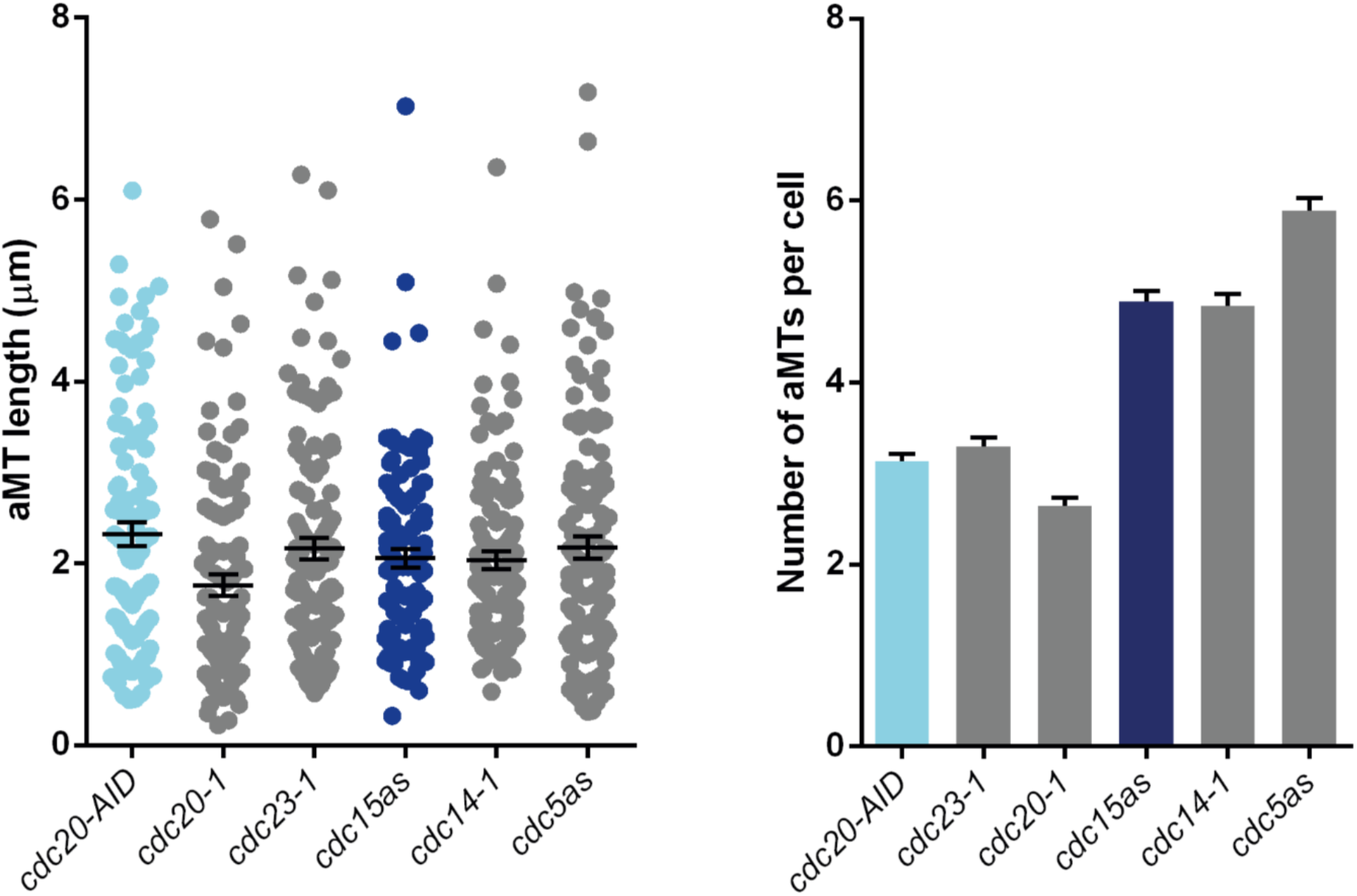
The aMT morphology is dictated by the cell cycle phase. *cdc20-AID* (Ry7873), *cdc20-1* (Ry586), *cdc23-1* (Ry454), *cdc14-1* (Ry1574), *cdc5-as1* (Ry2446), and *cdc15-as1* (Ry1112) cells were synchronized in G1 and released at 37°C, to inactivate the *cdc20-1*, *cdc23-1* and *cdc14-1* alleles, into fresh YEPD media supplemented with Auxin (500µM), 1NM-PP1 analogue 9 (5µM) and CMK (5µM) to inactivate the *cdc20-AID, cdc15-as1* and *cdc5-as1* alleles, respectively. Cells were analyzed at their terminal arrest (∼3 hours after the release). The graphs show the aMT length and number of the indicated mutant strains (n=100).

**Fig. S2.**
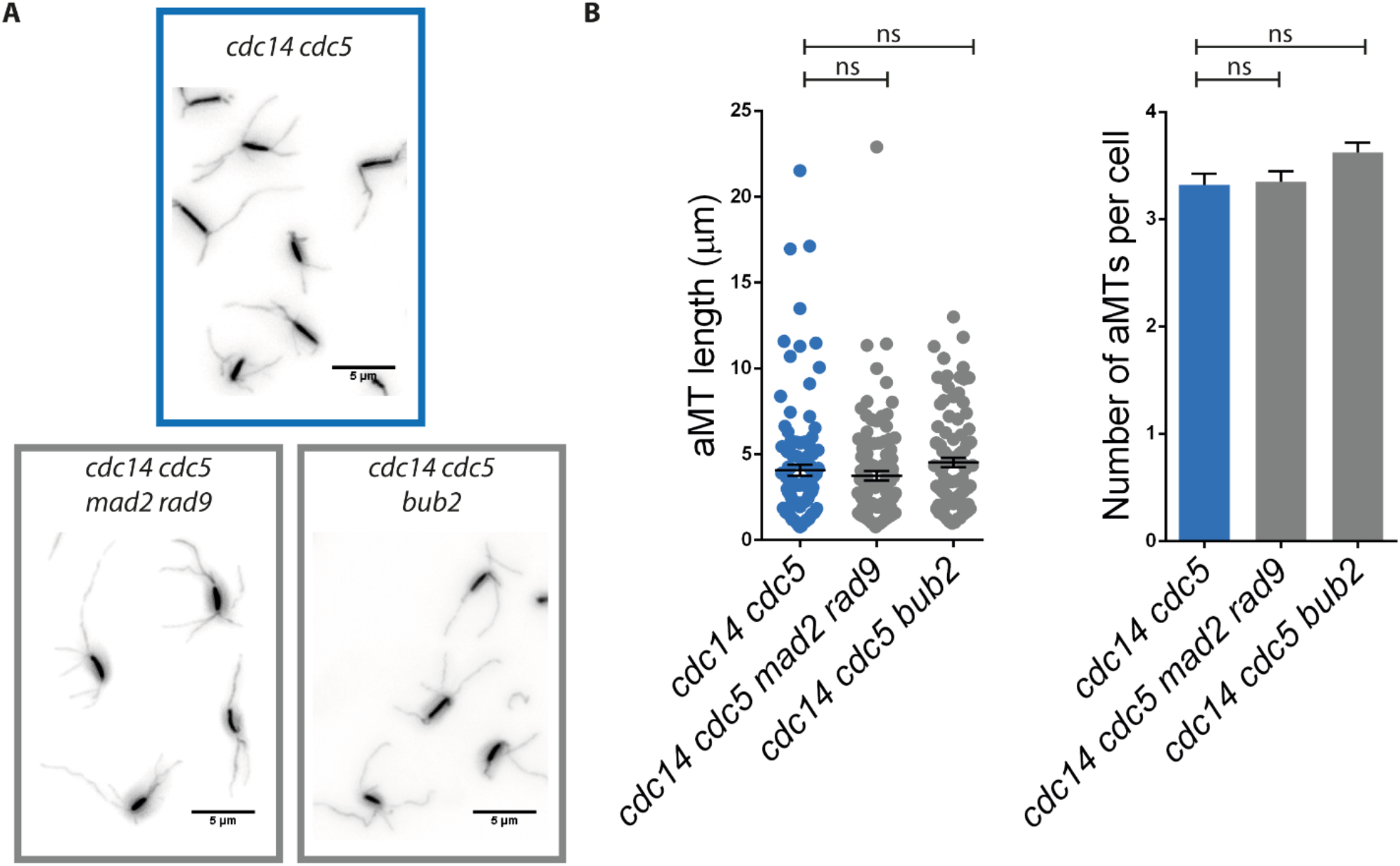
The aMT phenotype of *cdc14-1 cdc5-as1* cells is independent from activation of the SAC, the DDR and the SPoC. **(A-B)** *cdc14-1 cdc5-as1* (Ry1602)*, cdc14-1 cdc5-as1 mad2Δ rad9*Δ (Ry3771) and *cdc14-1 cdc5-as1 bub2*Δ (Ry3346) cells were arrested in G1 and released at 37°C, to inactivate the *cdc14-1* allele, into fresh YEPD media supplemented with CMK (5µM) to inactivate the *cdc5-as1* allele. Cells were scored at their terminal arrest (∼3 hours after the release). **(a)** Representative images of the indicated strains are shown (scale bar = 5µm). **(b)** The graphs show the aMT length and number of the indicated strains (n=100).

**Fig. S3.**
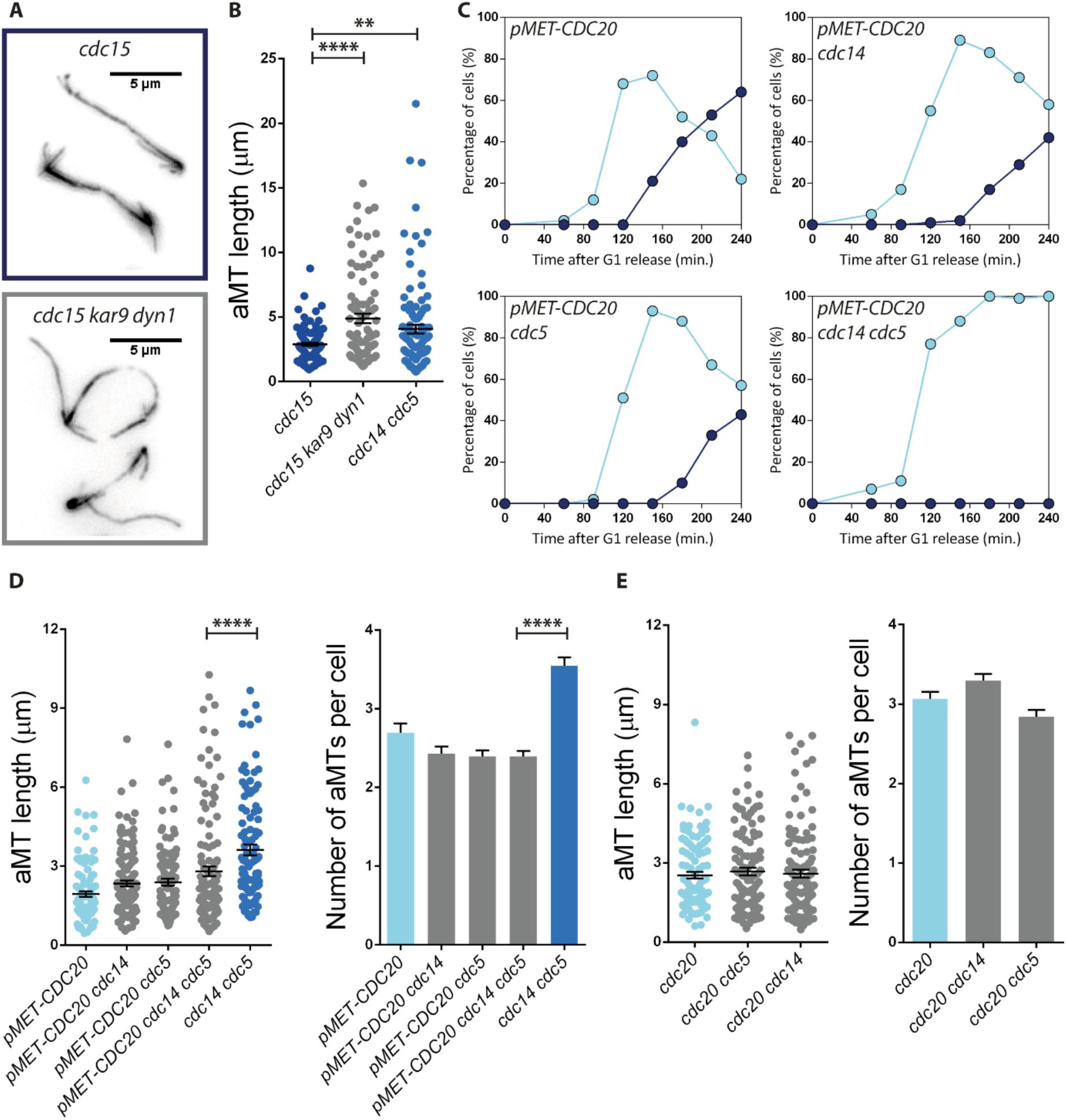
The aMT phenotype of *cdc14-1 cdc5-as1* cells is not a consequence of the spindle elongation defect or the inactivation of Cdc14 and Cdc5. **(A-B)** *cdc15-as1* (Ry1112), *cdc15-as1 kar9Δ dyn1-AID* (Ry7620), and *cdc14-1 cdc5-as1* (Ry1602) cells were arrested in G1 and released at 37°C, to inactivate the *cdc14-1* allele, into fresh YEPD, supplemented with Auxin (500µM), 1NM-PP1 analogue 9 (5µM) and CMK (5 µM) to inactivate the *dyn1-AID*, *cdc15-as1* and *cdc5-as1* alleles, respectively. Cells were analyzed at their terminal arrest (∼3 hours after the release). **(A)** Representative images of *cdc15-as1* and *cdc15-as1 kar9Δ dyn1-AID* mutant cells are shown (scale bar = 5µm). **(B)** aMT length and number are shown for n=100 cells *per* strain (**=p<0.01; ****=p<0.0001; * asterisks denote significant differences according to ordinary One-Way ANOVA and Tukey’s multiple comparisons test). **(C-D)** *pMET-CDC20* (Ry1223), *pMET-CDC20 cdc14-1* (Ry3204) *pMET-CDC20 cdc5-as1* (Ry3209), *pMET-CDC20 cdc14-1 cdc5-as1* (Ry3201), and *cdc14-1 cdc5-as1* (Ry1602) cells were arrested in G1 with the α-factor pheromone (5µg/ml) in synthetic complete media lacking methionine (SC-Met) and synchronously released into YEPD media supplemented with methionine - to repress the expression of *CDC20 -* at 37°C in the presence of CMK (5 µM) to inactivate the *cdc14-1* and *cdc5-as1* alleles, respectively. **(C)** Samples were taken at the indicated time points to determine the percentage of cells with metaphase-like (light blue) and anaphase-like (dark blue) spindles (n=100). **(D)** aMT length and number were scored 3,5 hours after the release (n=100; ****=p<0.0001; * asterisks denote significant differences according to ordinary One-Way ANOVA and Tukey’s multiple comparisons test). **(E)** *cdc20-AID* (Ry4853)*, cdc20-AID cdc14-1* (Ry4934) and *cdc20-AID cdc5-as1* (Ry4936) cells were arrested in G1, released into fresh YEPD media at 37°C to inactivate the *cdc14-1* allele, and supplemented with Auxin (500µM) and CMK (5µM) to inactivate the *cdc20-AID* and *cdc5-as1* alleles, respectively. aMT length and number were analyzed when cells reached the terminal arrest (∼3 hours after the release) (n=100).

**Fig. S4.**
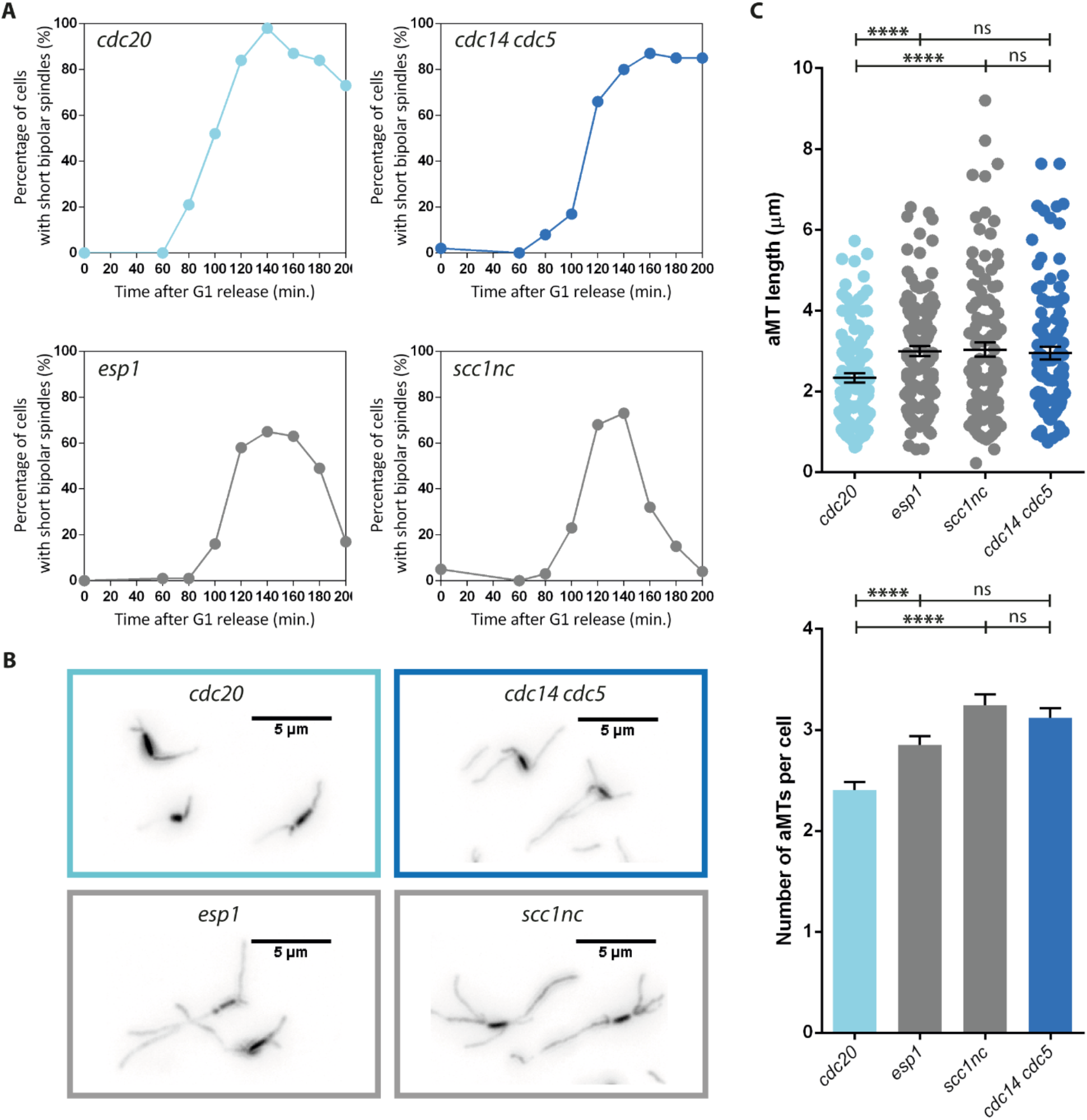
APC/C^Cdc20^ activation is sufficient to stabilize aMTs. **(A-C)** *cdc20-AID* (Ry4853), *esp1-1* (Ry9490), *pGAL-SCC1nc* (Ry8210), and *cdc14-1 cdc5-as1* (Ry1602) cells were synchronized in G1 in YEPR media at 23°C and released into fresh YEPR+G media, to express the *SCC1nc* allele, supplemented with Auxin (500µM) and CMK (5µM) to inactivate the *cdc20-AID* and *cdc5-as1* alleles, respectively. The cultures were incubated at 37°C to inactivate the *esp1-1* and *cdc14-1* alleles. **(A)** Samples were taken at the indicated time points to determine the percentage of cells with short bipolar spindles (n=100). **(B)** aMT length and number were scored 140 minutes after the release, the time point before spindle collapse, in *esp1-1* and *scc1nc* cells (n=100; ****=p<0.0001; * asterisks denote significant differences according to ordinary One-Way ANOVA and Tukey’s multiple comparisons test). **(C)** Representative images of *cdc20-AID*, *esp1-1*, *pGAL-SCC1nc*, and *cdc14-1 cdc5-as1* cells are shown (scale bar = 5µm).

**Fig. S5.**
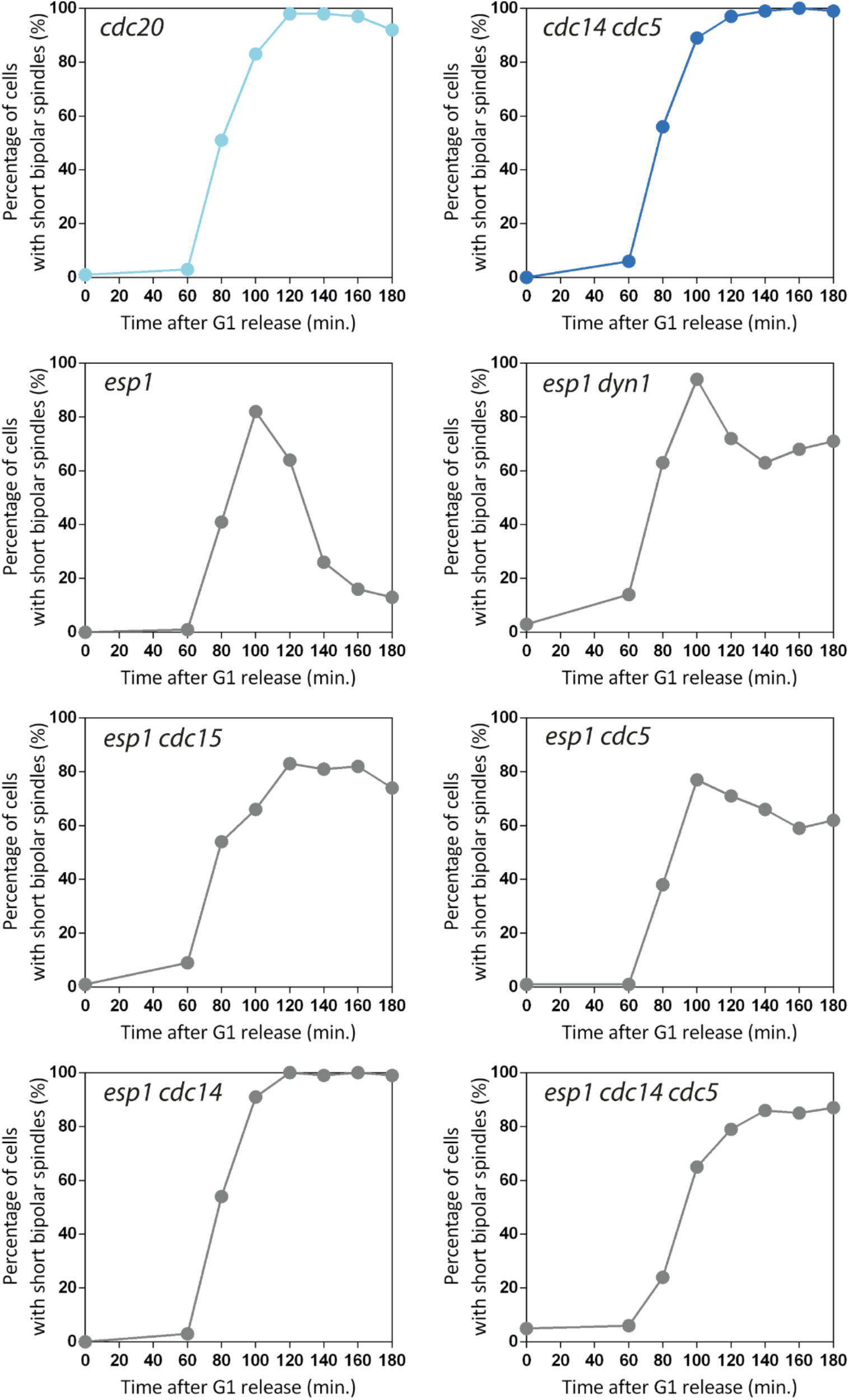
Preventing MEN activation or nuclear movement into the bud stops spindle collapse in *esp1-1* cells. *esp-1* (Ry9490), *esp1-1 dyn1*Δ (Ry9516), *esp1-1 cdc15-as1* (Ry9512), *esp1-1 cdc5-as1* (Ry9134*), esp1-1 cdc14-1* (Ry9131), *esp1-1 cdc14-1 cdc5-as1* (Ry9128), and *cdc14-1 cdc5-as1* (Ry1602) cells were arrested in G1 and released at 37°C, to inactivate the *esp1-1* and *cdc14-1* alleles, into fresh YEPD media supplemented with 1NM-PP1 analogue 9 (5µM) and CMK (5µM) to inactivate the *cdc15-as1* and *cdc5-as1* alleles, respectively. Samples were taken at the indicated time points to determine the percentage of cells with short bipolar spindles (n=100).

**Fig. S6.**
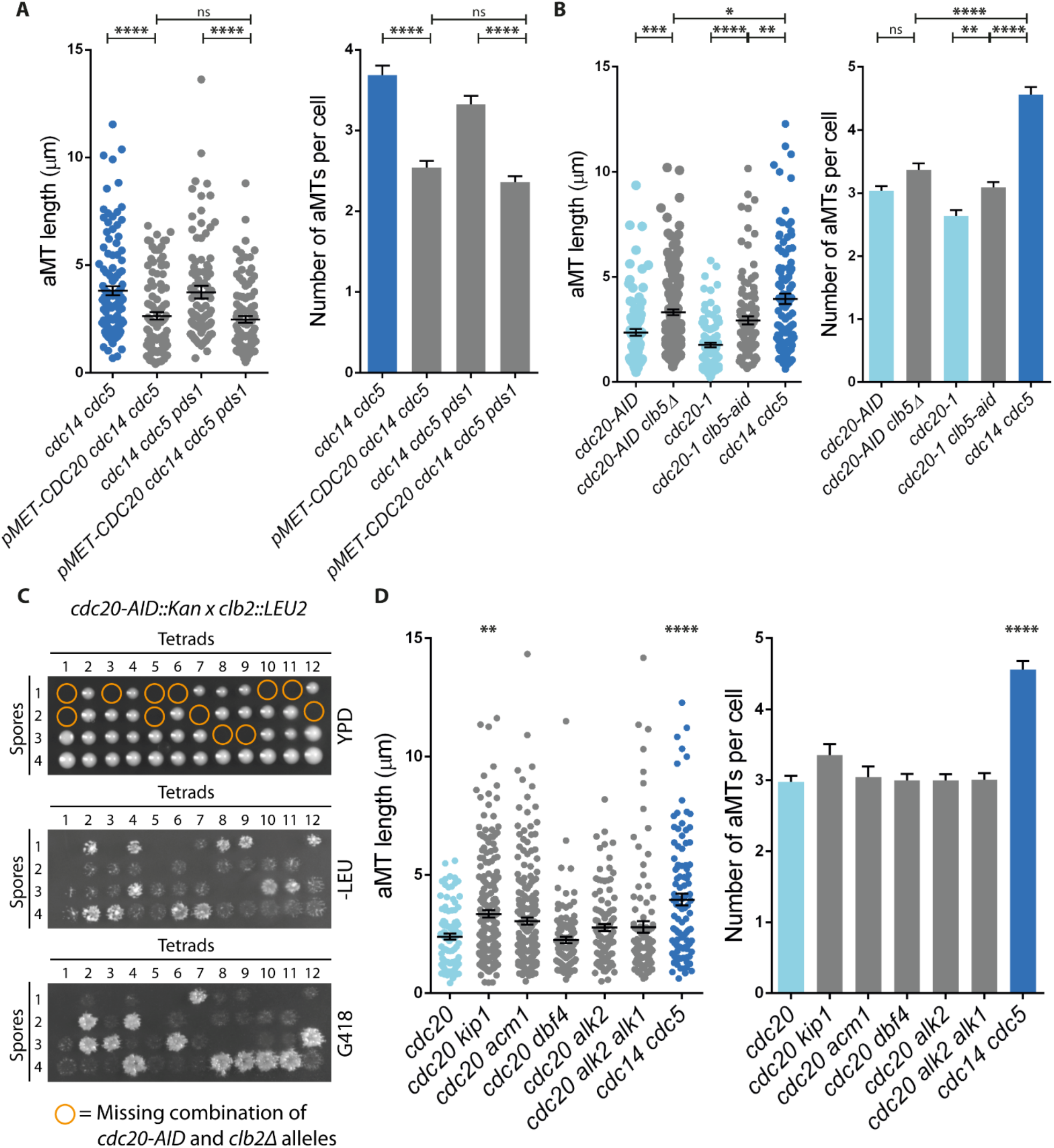
The individual removal of Pds1 and the majority of known or putative APC/C^Cdc20^ substrates does not alter aMT dynamics. **(A-B)** *cdc14-1 cdc5-as1* (Ry1602*), pMET-CDC20 cdc14-1 cdc5-as1* (Ry3201), *pds1Δ cdc14-1 cdc5-as1* (Ry2143), and *pMET-CDC20 pds1Δ cdc14-1 cdc5-as1* (Ry8969) cells were arrested in G1 with α-factor (5µg/ml) in synthetic complete media lacking methionine (SC-Met) and released into YEPD media lacking the pheromone and supplemented with methionine and CMK (5µM) to repress the expression of *CDC20* and to inactivate the *cdc5-as1* allele, respectively. The culture was incubated at 37°C to inactivate the *cdc14-1* allele. **(a)** aMT length and number were analyzed at the terminal arrest (∼ 3,5 hours after the release) (n=100; ****=p<0.0001; * asterisks denote significant differences according to ordinary One-Way ANOVA and Tukey’s multiple comparisons test). **(B)** *cdc20-AID* (Ry7873)*, cdc20-AID clb5*Δ (Ry9291)*, cdc20-1* (Ry586), *cdc20-1 clb5-AID* (Ry9760) and *cdc14-1 cdc5-as1* (Ry1602) cells were arrested in G1 and released into fresh YEPD media at 37°C to inactivate the *cdc20-1* and the *cdc14-1* alleles and supplemented with CMK (5µM) to inactivate the *cdc5-as1* allele. Auxin (500µM) was added (i) at the release to inactivate the *cdc20-AID* allele or (ii) when the majority (>90%) of the population reached metaphase (∼2 hours after the release) to inactivate the *clb5-AID* allele. Samples were taken 180 minutes after the release. aMT length and number are shown (n=100; *=p<0.1; **=p<0.01; ***=p<0.001; ****=p<0.0001; * asterisks denote significant differences according to ordinary One-Way ANOVA and Tukey’s multiple comparisons test). **(C)** Synthetic lethality of combining *cdc20-AID* and *clb2*Δ alleles seen by tetrad dissection. **(D)** *cdc20-AID* (Ry7873)*, cdc20-AID kip1*Δ (Ry9294)*, cdc20-AID acm1*Δ (Ry10025), *cdc20-AID dbf4-1* (Ry9877)*, cdc20-AID alk2*Δ (Ry9880)*, cdc20-AID alk2Δ alk1*Δ (Ry9883)*, cdc20-AID clb3*Δ (Ry10738), and *cdc14-1 cdc5-as1* (Ry1602) cells were arrested in G1 and released at 37°C, to inactivate the *cdc14-1* allele, into fresh YEPD media supplemented with Auxin (500µM) and CMK (5µM) to inactivate the *cdc20-AID* and *cdc5-as1* alleles, respectively. *cdc20-AID dbf4-1* cells were treated as the other strains, but incubated at 37°C when the majority of the cells reached metaphase (∼2 hours after the release). aMT length and number were scored at the terminal arrest (n=100; **=p<0.01; ****=p<0.0001; * asterisks denote significant differences according to ordinary One-Way ANOVA and Tukey’s multiple comparisons test against the control strain *cdc20-AID*).

**Fig. S7.**
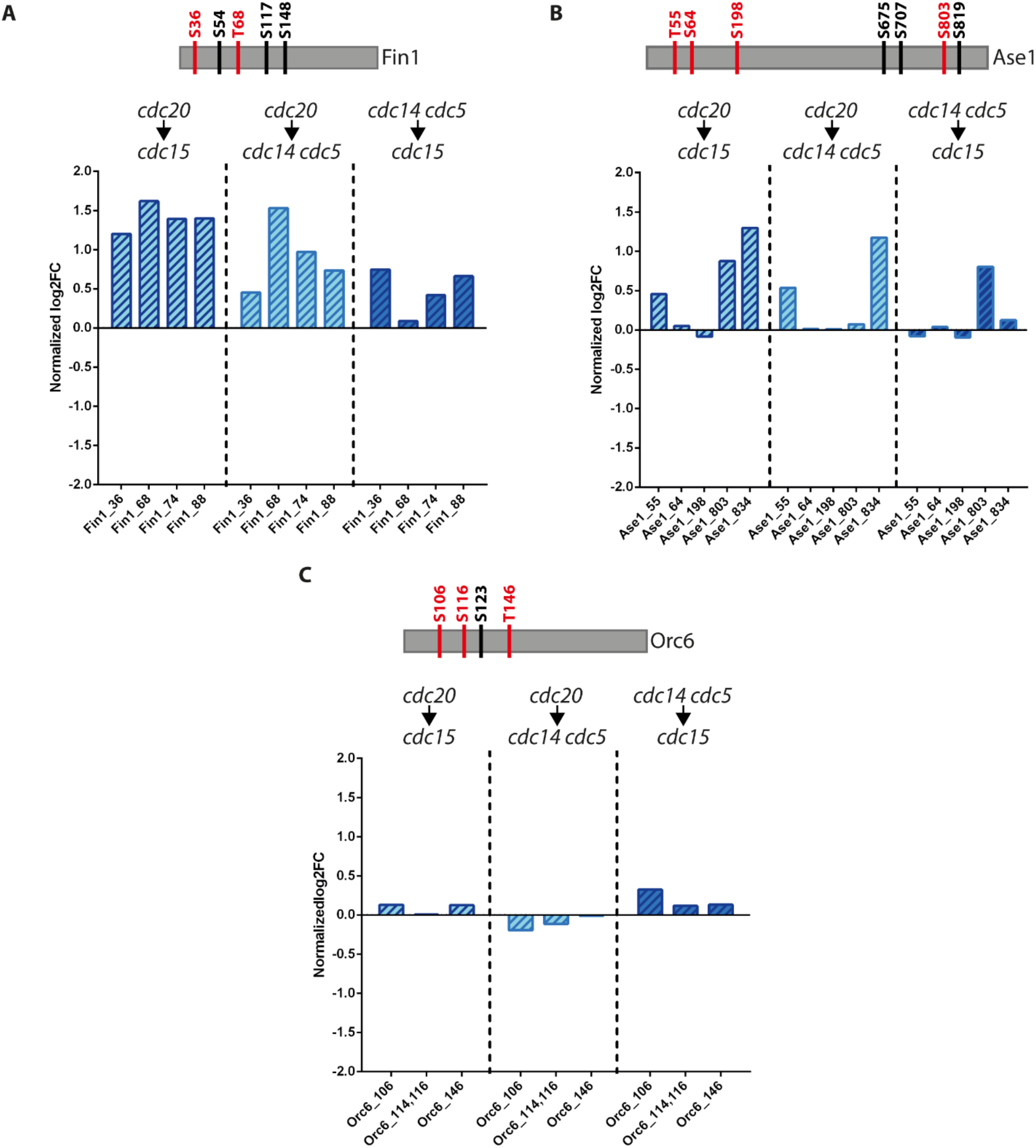
The de-phosphorylation kinetic of different CDK substrates is recapitulated by the phospho-proteomic analysis of *cdc20-aid*, *cdc14-1 cdc5-as1* and *cdc15-as1* cells. As a proof of concept that our phospho-proteomic analysis is a good proxy for cell cycle progression, we hand-picked and scored the phosphorylation status of residues in three CDK substrates, namely Fin1 **(A)**, Ase1 **(B)** and Orc6 **(C)**, whose kinetics of de-phosphorylation differs in anaphase. More precisely, these substrates are de-phosphorylated in early (Fin1), mid (Ase1) and late anaphase (Orc6). Above each graph, a schematic representation shows the identified putative CDK-phosphorylated residues covered by our analysis (highlighted in red), and the residues that were not identified in the data-set (highlighted in black). The graphs show the log2 fold change of each residue generated comparing each mutant with each other. The log2 fold changes of each residue are normalized by the log2 fold changes of the respective proteins to account for changes in protein abundance.

**Table S1.** List of strains used in this study

| Strain (Ry) | Relevant genotype | Origin |
| --- | --- | --- |
| 1 | *MATa, ade2-1, leu2-3, ura3, trp1-1, his3-11,15, can1-100, GAL, psi+* | Visintin lab |
| 20 | *MATa, clb2::LEU2* | Visintin lab |
| 586 | *MATa,cdc20-1* | Visintin lab |
| 1112 | *MATa, cdc15::CDC15-as1(L99G)::URA3* | Visintin lab |
| 1223 | *MATa, pMET3-CDC20::URA3, CDC14-3HA* | Visintin lab |
| 1574 | *MATa, cdc14-1* | Visintin lab |
| 1602 | *MATa, cdc14-1, cdc5-as1(L158G)* | Visintin lab |
| 2143 | *MATa, cdc14-1, cdc5-as1(L158G), pds1::URA3* | Visintin lab |
| 2446 | *MATa, cdc5-as1(L158G)* | Visintin lab |
| 3201 | *MATa, pMET3-CDC20::URA3, cdc14-1, cdc5-as1(L158G)* | Visintin lab |
| 3204 | *MATa, pMET3-CDC20::URA3, cdc14-1* | Visintin lab |
| 3209 | *MATa, pMET3-CDC20::URA3, cdc5-as1(L158G), CDC14-3HA* | Visintin lab |
| 3256 | *MATa, cdc14-1, cdc5-as1(L158G), ura3::pAFS125-TUB1p-GFPTUB1::URA3* | Visintin lab |
| 3346 | *MATa, cdc14-1, cdc5-as1(L158G), bub2::HIS3* | Visintin lab |
| 3771 | *MATa, cdc14-1, cdc5-as1(L158G), mad2::URA3, rad9::LEU2* | Visintin lab |
| 4853 | *MATa, ura3::pADH1-OsTIR1-9MYC::URA3, CDC20-aid::KanMX* | Visintin lab |
| 4854 | *MATalpha, ura3::pADH1-OsTIR1-9MYC::URA3, CDC20-aid::KanMX* | Visintin lab |
| 4936 | *MATa, cdc5-as1(L158G), ura3::pADH1-OsTIR1-9MYC::URA3, CDC20-aid::KanMX* | Visintin lab |
| 7545 | *MATa, kar9::HIS5, ura3::pADH1-OsTIR1-9MYC::URA3, DYN1-aid::KanMX, CDC20-aid::KanMX* | This Manuscript |
| 7589 | *MATa, cdc5-as1(L158G), kar9::HIS5* | This Manuscript |
| 7620 | *MATa, leu2::pTEF1-osTIR::LEU2, DYN1-aid::KanMX, kar9::HIS5, cdc15-as1(L99G)::URA3* | This Manuscript |
| 7623 | *MATa, leu2::pTEF1-osTIR::LEU2, DYN1-aid::KanMX, cdc15-as1(L99G)::URA3* | This Manuscript |
| 7626 | *MATa, kar9::HIS5, cdc15-as1(L99G)::URA3* | This Manuscript |
| 7694 | *MATa, leu2::pTEF1-osTIR::LEU2, DYN1-aid:KanMX, kar9::HIS5, cdc5-as1(L158G)* | This Manuscript |
| 7697 | *MATa, leu2::pTEF1-osTIR::LEU2, DYN1-aid:KanMX, cdc5-as1(L158G)* | This Manuscript |
| 7702 | *MATa, cdc5-as1(L158G), leu2::pTEF1-osTIR::LEU2, kar9::HIS5, CDC20-aid::KanMX* | This Manuscript |
| 7732 | *MATa, leu2::pTEF1-osTIR::LEU2, CDC20-aid::KanMX, ura3::pAFS123-TUB1p-GFPTUB1::URA3* | This Manuscript |
| 7746 | *MATa, leu2::pTEF1-osTIR::LEU2, DYN1-aid:KanMX, kar9::HIS5, CDC20-aid::KanMX, cdc5-as1(L158G)* | This Manuscript |
| 7749 | *MATa, leu2::pTEF1-osTIR::LEU2, DYN1-aid:KanMX, CDC20-aid::KanMX, cdc5-as1(L158G)* | This Manuscript |
| 7873 | *MATa, leu2::pTEF1-osTIR::LEU2, CDC20-aid::KanMX* | This Manuscript |
| 8210 | *MATa, pGAL-SCC1(R180D,R268D)* | This Manuscript |
| 8969 | *MATa, pMET3-CDC20::URA3, cdc14-1, cdc5-as1(L158G), pds1::URA3* | This Manuscript |
| 9128 | *MATa, cdc14-1, cdc5-as1(L158G), esp1-1* | This Manuscript |
| 9131 | *MATa, cdc14-1, esp1-1* | This Manuscript |
| 9134 | *MATa, cdc5-as1(L158G), esp1-1* | This Manuscript |
| 9237 | *MATa, ura3::pADH1-OsTIR1-9MYC::URA3, CDC20-aid::KanMX, cdc12-6* | This Manuscript |
| 9291 | *MATa, leu2::pTEF1-osTIR::LEU2, CDC20-aid::KanMX, clb5::URA3* | This Manuscript |
| 9294 | *MATa, leu2::pTEF1-osTIR::LEU2, CDC20-aid::KanMX, kip1::HIS3* | This Manuscript |
| 9490 | *MATa, esp1-1* | This Manuscript |
| 9512 | *MATa, esp1-1, cdc15-as1(L99G)::URA3* | This Manuscript |
| 9516 | *MATa, leu2::pTEF1-osTIR::LEU2, CDC20-aid::KanMX, cdc15-as1(L99G)::URA3* | This Manuscript |
| 9741 | *MATa, cdc15::CDC15-as1(L99G)::URA3, ura3::pAFS125-TUB1p-GFPTUB1::URA3* | This Manuscript |
| 9760 | *MATa, cdc20-1, CLB5-aid:KanMX* | This Manuscript |
| 9877 | *MATa, leu2::pTEF1-osTIR::LEU2, CDC20-aid::KanMX, dbf4-1* | This Manuscript |
| 9880 | *MATa, leu2::pTEF1-osTIR::LEU2, CDC20-aid::KanMX, alk2::HIS3* | This Manuscript |
| 9883 | *MATa, leu2::pTEF1-osTIR::LEU2, CDC20-aid::KanMX, alk2::HIS3, alk1::KanMX* | This Manuscript |
| 10025 | *MATa, leu2::pTEF1-osTIR::LEU2, CDC20-aid::KanMX, acm1::KanMX* | This Manuscript |
| 10741 | *MATa, leu2::pTEF1-osTIR::LEU2, CDC20-aid::KanMX, clb4::HIS3* | This Manuscript |
| 10844 | *MATa, cdc14-1, cdc5-as1(L158G), kip2::HphMX6* | This Manuscript |
| 10857 | *MATa, cdc14-1, cdc5-as1(L158G), kip2::HphMX6, pCEN-KIP2-13myc(\*)* | This Manuscript |
| 10858 | *MATa, leu2::pTEF1-osTIR::LEU2, CDC20-aid::KanMX, kip2::HphMX6, pCEN-KIP2-13myc(\*)* | This Manuscript |
| 10861 | *MATa, leu2::pTEF1-osTIR::LEU2, CDC20-aid::KanMX, kip2::HphMX6, pCEN-KIP2-S63A-T275A-13myc(\*)* | This Manuscript |

(\*) Plasmids received from Liakopoulos lab. (Drechsler et al., 2015)

**Other Supplementary Materials for this manuscript include the following:**

**Table S2**

**Table S3**

